# MolAI: A Deep Learning Framework for Data-driven Molecular Descriptor Generation and Advanced Drug Discovery Applications

**DOI:** 10.1101/2025.03.24.644888

**Authors:** Sayyed Jalil Mahdizadeh, Leif A. Eriksson

**Affiliations:** Department of Chemistry and Molecular Biology, University of Gothenburg, 405 30 Göteborg, Sweden

**Keywords:** Latent space, Molecular representation, Drug discovery, Autoencoder NMT, Molecular descriptors

## Abstract

This study introduces MolAI, a robust deep learning model designed for data-driven molecular descriptor generation. Utilizing a vast training dataset of 221 million unique compounds, MolAI employs an autoencoder neural machine translation (NMT) model to generate latent space representations of molecules. The model demonstrated exceptional performance through extensive validation, achieving a 99.99% accuracy in regenerating input molecules from their corresponding latent space. This study showcases the effectiveness of MolAI-driven molecular descriptors by developing an ML-based model (iLP) that accurately predicts the predominant protonation state of molecules at neutral pH. These descriptors also significantly enhance ligand-based virtual screening and are successfully applied in a framework (iADMET) for predicting ADMET features with high accuracy. This capability of encoding and decoding molecules to and from latent space opens unique opportunities in drug discovery, structure-activity relationship analysis, hit optimization, *de novo* molecular generation, and the training infinite machine learning models.

**Graphical Abstract:** 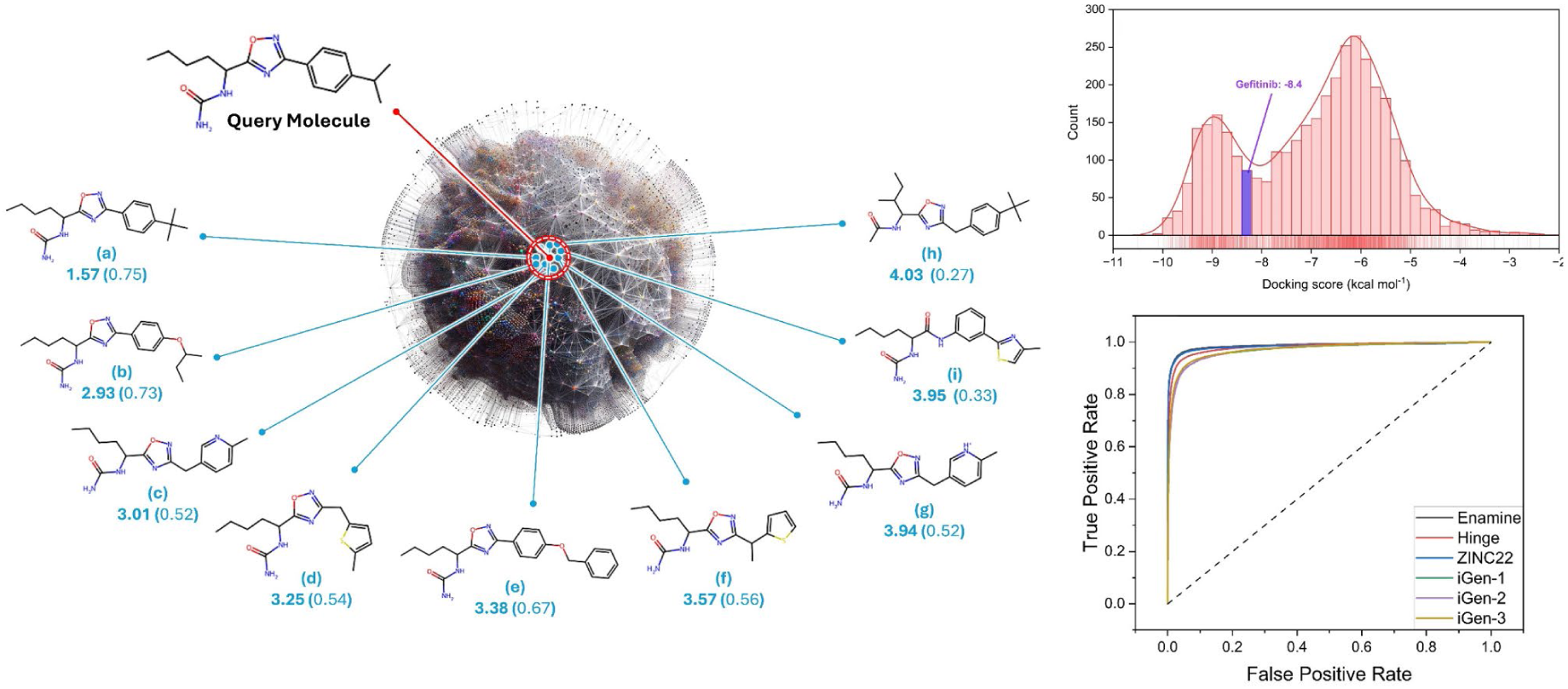

## 1. INTRODUCTION

Recent breakthroughs in artificial intelligence (AI) and machine learning (ML) have revolutionized the field of cheminformatics (*1, 2*) and drug discovery (*3–5*), enabling the prediction of molecular properties, the design of novel compounds with desired characteristics, and the estimation of drug/bio-macromolecular interactions. The core of these advancements lies in the development of molecular descriptors, which are quantitative representations of chemical information of actual molecules used in various predictive and generative models. Traditionally, these descriptors were crafted manually, relying on expert knowledge to encode molecular properties into a computer-interpretable vector (*6*). The most employed vector of molecular representations is Morgan fingerprints, also known as extended-connectivity fingerprints (ECFPs), as these often outperform other types of fingerprints in molecular bioinformatics and virtual screening tasks (*7, 8*). However, ECFPs are not only very high dimensional and sparse but may also suffer from bit collisions introduced by the hashing step (*9*) On the other hand, most of the ML models developed in the domain of cheminformatics and drug design commonly employ pre-extracted traditional molecular descriptors as input which contrasts with the core principle of representation learning (*10*). In particular, in deep learning (*11*) a robust data representation must be driven from a simple yet complete featurization, rather than depending on human-engineered descriptors.

The advancements in deep learning algorithms have paved the way for data-driven approaches that automatically learn molecular representations from large datasets of lower-level molecular formats such as molecular graphs (*12*) or SMILES (simplified molecular input line entry specification) (*13*). In this context, 2018 was a landmark year, with the publication of three sophisticated studies that significantly advanced the field. Jaeger *et al.* (*9*) introduced mol2vec, an unsupervised ML approach with chemical Intuition, an innovative method inspired by natural language processing (NLP) techniques, to create vector representations of molecular substructures. This approach leverages the principles of word2vec(*14*), which learns high-dimensional embeddings of words based on their context in sentences. Similarly, mol2vec treats molecular substructures as “words” and entire molecules as “sentences.” mol2vec utilizes the Morgan algorithm to generate unique identifiers for substructures within a molecule, which are then ordered to form “molecular sentences”. These sentences are processed using word2vec to obtain dense vector representations of the substructures. The resulting vectors can be summed to represent entire molecules, which can then be used as features in supervised machine learning tasks to predict molecular properties and activities.

Gómez-Bombarelli *et al.* (*15*) explored a novel approach to molecular design utilizing a variational autoencoder (VAE) (*16*) to transform discrete SMILES molecular representations to and from a continuous multidimensional vector. They demonstrated that this continuous representation allows for the automatic generation of novel chemical structures and could also be used as descriptors for down-stream ML-based prediction tasks. The autoencoder is composed of two neural networks: the encoder and the decoder. The encoder (based on the 1D convolutional neural network (CNN) (*17*)) transforms a variable-length SMILES sequence into a fixed-size continuous latent representation (latent space). The decoder network (based on the gated recurrent unit (GRU) (*18*)) then takes this latent space and reconstructs it back into the original input sequence (seq2seq learning (*19*)). The training objective for the entire autoencoder is to minimize the mean reconstruction error at the character level for each input sequence. The design includes an information bottleneck between the encoder and decoder, which compels the autoencoder network to condense the critical information into the latent space. This ensures that the decoder can accurately reconstruct the input sequence while retaining as much information as possible.

Winter *et al.* (*20*) however, argued that training an autoencoder to reconstruct a sequence representing a molecule poses the risk that the network may primarily learn syntactic features and repetitive patterns within the sequence. This focus can lead to the neglect of semantic information, such as molecular properties, resulting in a failure to encode higher-level features that are crucial for accurate molecular representation. Consequently, the network might capture superficial patterns instead of the essential chemical and structural characteristics needed for meaningful molecular analysis and prediction. They therefore addressed this challenge by proposing a method based on translation rather than reconstruction. This approach is inspired by the process of human translation, where a person reads an entire sentence to grasp its meaning before translating it into another language. Similarly, a neural machine translation (NMT) (*21*) model processes the entire input sequence and encodes it into an intermediate continuous vector representation, known as the latent space. This latent space encapsulates the “understanding” of the input sequence’s meaning, including all relevant semantic information shared by the input and output sequences. Their NMT autoencoder model consists of stacked 3×GRU units for both encoder and decoder parts with a dense “bottleneck” layer of length 512 between encoder and decoder as the latent vector.

Inspired by Winter *et al.*(*20*), the current study adopts a similar NMT approach to train an autoencoder model for data-driven molecular descriptors, while addressing certain limitations and incorporating improvements:

- The size of the training dataset has been enlarged by more than 200% to enhance robustness, generalization, diversity, and to reduce the risk of overfitting.
- To further augment the autoencoder model in learning a meaningful molecular representation in terms of physiochemical features, we reinforced the translation model by incorporating regression models for 13 molecular properties, compared to 9 in the original model.
- GRU units, utilized in the original translation model, perform well on low-complexity sequences while for high-complexity sequences, long short-term memory (LSTM)(*22*) units are typically better at capturing long-term dependencies due to its more complex architecture, thereby outperforming GRUs (*23*).
- The training and prediction speed has been significantly accelerated by using TensorFlow 2.7 instead of TensorFlow 1.1 in the original model. TensorFlow 2.7 represents a profound evolution from TensorFlow 1.1, focusing on ease of use, flexibility, and performance (www.tensorflow.org/guide/migrate/tf1_vs_tf2).

By addressing these points, we aim to build a more robust, generalizable, and efficient model, named “MolAI”, for data-driven molecular descriptors that could be used for training an infinite number of down-stream molecular ML-based models. The quality and applicability of the resulting data-driven molecular descriptors have been comprehensively evaluated through various benchmarking experiments. These experiments range from identifying the predominant protonation state of molecules in aqueous solutions at a given pH (referred to as iLP), to *de novo* molecular generation, improving the enrichment factor (EF) in ligand-based virtual screening, and predicting ADMET (Absorption, Distribution, Metabolism, Excretion, and Toxicity) features of drug-like molecules (referred to as iADMET). The precise protonation state of molecules is critically important in molecular docking, virtual screening, and drug discovery because it not only directly influences the accuracy of computational predictions (*24–26*) but also have a significant effect on the screening time (*27, 28*). Protonation states significantly affect the charge distribution and hydrogen bonding patterns of molecules (*29*), which are essential for correctly modeling molecular interactions. Accurate protonation states ensure that the geometric and electronic properties of both ligands and target proteins are appropriately represented, leading to more reliable docking results. Inaccurate protonation states lead to accumulated false positives or false negatives, reducing the efficiency of the screening process, and increasing the costs. The EF is a crucial metric in virtual screening that measures the effectiveness of a screening method in identifying active compounds from a vast library of decoys. It quantifies how much better a screening method is at selecting active compounds compared to random selection (EF=1). A high EF indicates that the method significantly enriches active compounds at the top of the ranked list, which is essential for efficient drug discovery. This metric is particularly important as it helps prioritize compounds for further experimental validation, thereby saving time and resources. The assessment of ADMET features is essential in drug discovery attempts (*30*) because it determines the pharmacokinetic and safety profile of potential drug candidates(*31*). Evaluating ADMET properties early in the drug development process helps in identifying compounds with favorable absorption and distribution characteristics, ensuring they reach therapeutic concentrations at the target site. It also assesses the metabolic stability and potential for drug-drug interactions, which are critical for predicting efficacy and avoiding adverse effects. Excretion profiles help understanding how the drug is eliminated from the body, impacting dosing and potential accumulation. Toxicity assessment is vital to ensure that compounds are safe for human use, minimizing the risk of harmful side effects.

## 2. RESULTS AND DISCUSSION

### 2.1. MolAI pretraining

Initially, the model architecture design and hyperparameter optimization was carried out using a smaller subset of 20 million randomly selected samples from the full training set of 221 million. The subset was further divided into training and validation sets (10:2). The Keras tuner tool was used for an extensive grid search based on the validation loss. This process involved monitoring the reduction in validation loss when varying the number (1, 2, 3) and size (256, 512, 1024) of the LSTM cells, learning rate (0.005, 0.001, 0.0001), batch size (64, 128, 256, 512), and latent vector dimension (256, 512, 1024). Over 300 models were investigated, each with a different combination of the parameters mentioned above, and the final model was chosen according to the combination of the hyperparameters resulting in the lowest validation loss as described in the Method section.

Figure 1a shows the loss (*categorical cross entropy*) and accuracy plots of the decoder part of the MolAI model after 50000 training steps. Figure 1b illustrates the loss function (*mean squared error*) of the regression models associated with prediction of 13 molecular properties, after 1000 training steps. As figures 1a and 1b indicate, a monotonical and smooth loss reduction was observed for the decoder and all regression prediction models implying that a tuneful set of hyperparameters led to an effective and well-balanced training process.

**Fig. 1.**
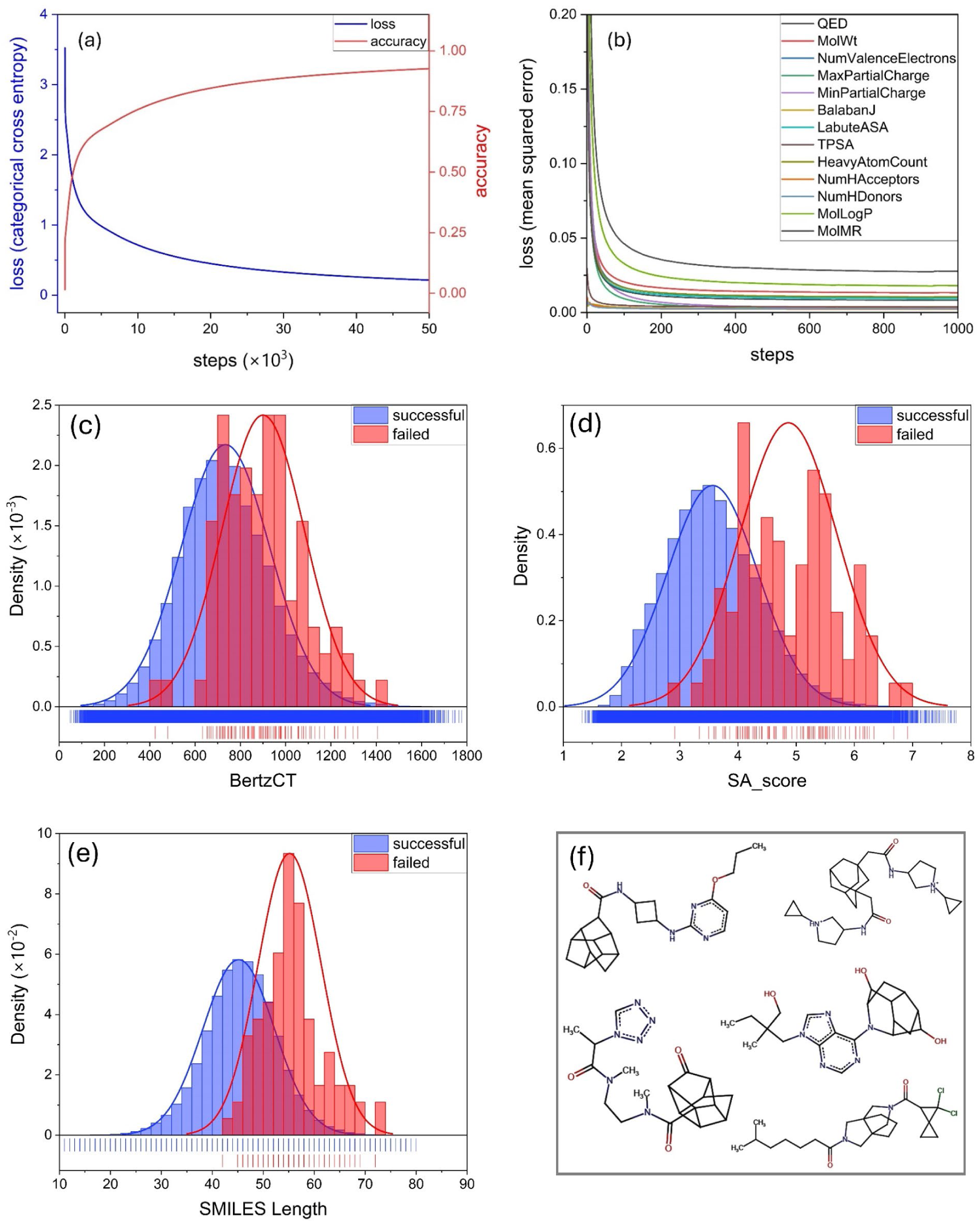
**(a)** Loss (categorical cross entropy) and accuracy plots of the decoder part of the MolAI model after 50000 training steps. **(b)** Loss function (mean squared error) of the regression models associated with prediction of 13 molecular properties, after 1000 training steps. The distribution of **(c)** Bertz complexity (BertzCT) index, **(d)** synthetic accessibility scores (SA_score), and **€** length of the SMILES strings for successful (blue) and failed (red) molecules during MolAI validation analysis. **(f)** 2D structures of five failed molecules were MolAI was not able to correctly reproduce the input molecules from the corresponding latent vectors.

### 2.2. MolAI validation

A set of 1 million compounds, randomly sampled from the ZINC22 database, was used to validate the model. Duplicates and overlaps with the MolAI training set were first removed, and the pre-processing steps (as detailed in the Method section) were conducted, resulting in 999917 unique molecules. The molecules were then tokenized and vectorized into a one-hot encoded format. The latent space vectors of the compounds were subsequently generated, and the accuracy of the MolAI autoencoder was assessed based on the success rate of regenerating the exact SMILES strings from the corresponding latent vectors. The validation task demonstrated near-perfect performance of MolAI, achieving an extremely high accuracy value of > 99.99% (only 91 failures) in correctly regenerating the input SMILES strings from the latent space vectors. Moreover, in 93.4% of the failed SMILES (85 out of the 91), the discrepancy between the original and the regenerated SMILES strings was limited to a single character.

Further investigation of the failed molecules did not reveal any particular pattern or presence of specific functional groups. However, distinct deviations in certain properties and metrics between the successful and failed molecules were observed (Figure 1). Figure 1c shows the distribution of Bertz complexity (BertzCT) index of successful (in blue) and failed (red) molecules. The BertzCT index is a topological measure that quantifies the “complexity” of molecules, consisting of two terms: one representing the complexity of bonding and the other representing the complexity of the distribution of heteroatoms (*32*). As indicated in Figure 1c, the distribution of the BertzCT index for the failed molecules is shifted significantly towards higher values, with a mean value of 900 compared to 734 for the successful molecules. This suggests that failed molecules exhibit higher structural complexity. A similar trend was observed for the distribution of synthetic accessibility scores (SA_score) as shown in Figure 1d. The SA_score, €d by Ertl and Schuffenhauer (*33*) estimates the ease of synthesis of organic molecules (ranging from 1 to 10, where 1 indicates very easy and 10 indicates very difficult to synthesize) based on a combination of fragment contributions and a complexity penalty. The average SA_score values for successful and failed molecules are 3.5 and 4.9, respectively, indicating that failed molecules are more challenging to synthesize. Additionally, the difference between successful and failed molecules is reflected in the length of the associated SMILES strings (Figure 1d). The average SMILES length of successful molecules is 45.1, significantly shorter than the 57.1 average length of the failed molecules. Figure 1f displays the 2D structures of several failed molecules, which predominantly feature multiple fused rings and cage-like substructures, further highlighting their high complexity.

### 2.3. Latent space evaluation

The continuous latent space enables quantitative similarity measurements by calculating the Euclidean distance between two molecules, treating them as distinct points in a high-dimensional space. This allows for a fast and efficient similarity search by sorting the Euclidean distances in ascending order between a query molecule and all molecules in a molecular library, as a lower Euclidean distance is indicative of higher similarity. Figure 2a illustrates an example of how the latent space and Euclidean distance between molecules could be employed for a similarity search. In Figure 2a, the number below each molecule indicates the Euclidean distance to the query molecule and the number in parentheses shows the Tanimoto similarity index (*34*) calculated using Morgan fingerprints as a vector of length 2048. The top two most similar compounds to the query molecule identified by MolAI and the Tanimoto index agree (molecules *a* and *b* in Figure 2a). However, the ranking of similar compounds diverges significantly thereafter. For example, the 3^rd^, 4^th^, 8^th^, and 9^th^ ranked molecules using MolAI (molecules *c*, *d*, *h*, and *i* in Figure 2a) were ranked 11^th^, 8^th^, 69^th^, and 268^th^ by the Tanimoto index, respectively. In addition, 23 (704) compounds were found with the same Tanimoto similarity index among the top 100 (top 1000) similar molecules, respectively, whereas MolAI provides a unique Euclidean distance for each compound. Moreover, MolAI can distinguish between different protonation states; for example, molecules *c* and *g* are two protonation states of the same molecule, showing the same Tanimoto index (0.52) but different Euclidean distances (3.01 and 3.94, respectively).

**Fig. 2.**
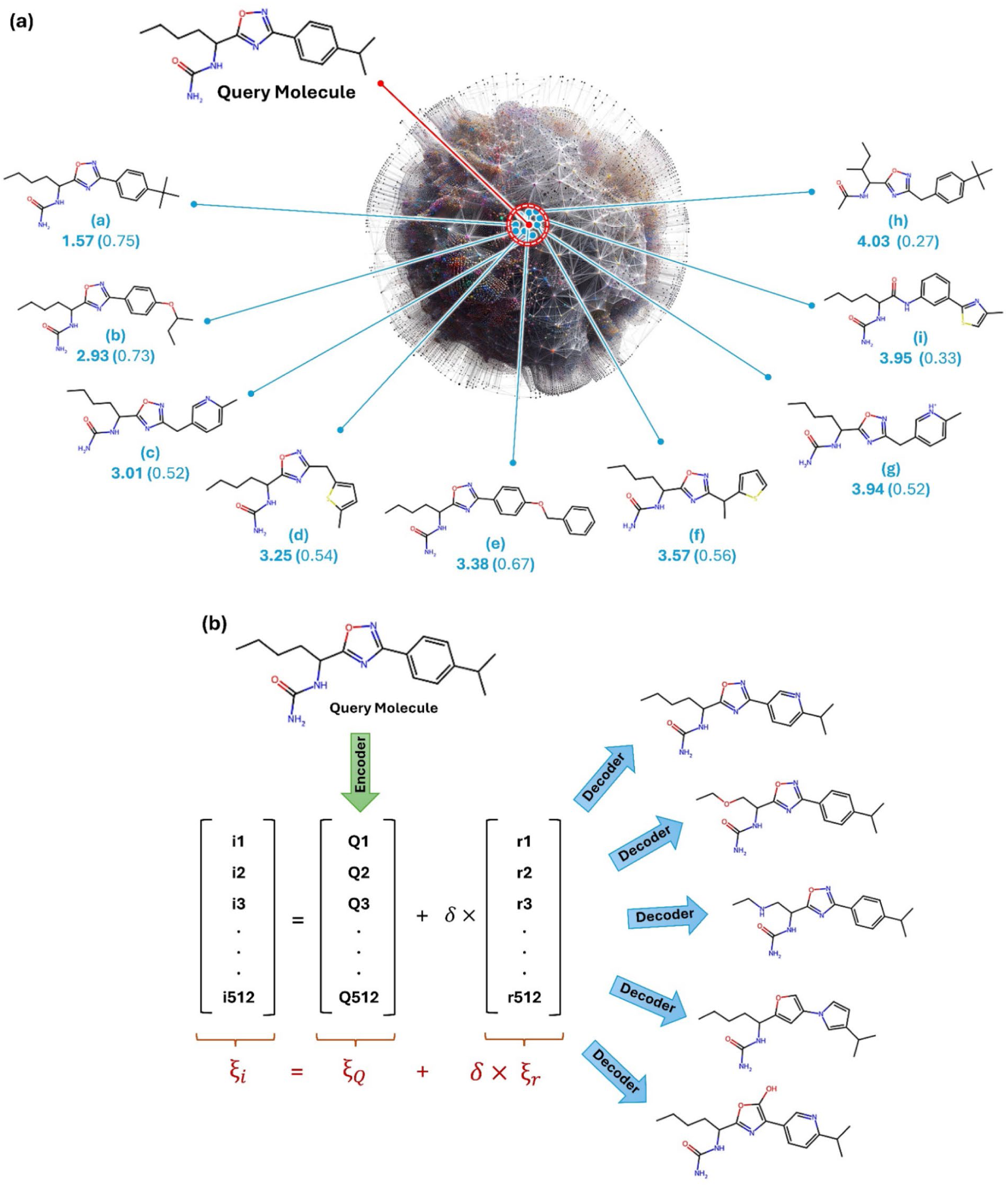
**(a)** An example of how the latent space and Euclidean distance between molecules can be employed in a similarity search. The number below each molecule indicates the Euclidean distance to the query molecule and the number in parentheses shows the Tanimoto similarity index. **(b)** Novel similar molecules and derivatives can be generated by creating a new latent vector (*ξ_i_*), by adding a random vector of length 512 (*ξ_r_*) generated using a standard normal distribution multiplied by a scaling factor *δ*, to the latent vector of the query molecule (*ξ_Q_*).

The differences in the similarity search results can be attributed to the distinct nature of the representations and similarity measures used. The MolAI autoencoder approach captures more complex and nuanced relationships between molecules, potentially identifying similarities that are relevant not only structurally but also physiochemically. In contrast, the Tanimoto similarity measure using Morgan fingerprints offers a simpler, substructure-based similarity assessment, which may overlook some of the subtler relationships that MolAI is able to capture.

Another advantage of a valid continuous latent space representation of molecular systems is the ability to generate similar molecules or derivatives by sampling the latent space around the latent vector of a query molecule, as illustrated in Figure 2b. Technically, a new latent vector (*ξ_i_*) is created by adding a random vector of length 512 (*ξ_r_*) generated using a standard normal distribution (with a mean of zero and standard deviation of one), multiplied by a scaling factor δ, to the latent vector of the query molecule (*ξ_Q_*):

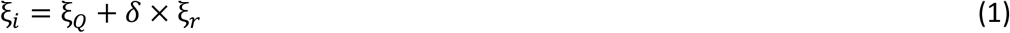

This process effectively perturbs the query molecule’s latent vector in a controlled manner, allowing for the generation of novel, yet similar, molecules. The decoder part of MolAI translates the perturbed latent vectors into the corresponding molecules. The scaling factor (*δ*) can be adjusted to control the degree of similarity between the generated molecules and the query molecule. A smaller *δ* results in molecules that are more similar to the query molecule, while a larger δ leads to more significant deviations, producing molecules that are structurally and potentially functionally different. However, depending on the density of the query latent vector’s locality in the latent space, this approach may result in many chemically invalid SMILES strings. This occurs because the latent space may contain regions that do not correspond to valid chemical structures, especially in sparsely populated areas. Tuning the scaling factor will be beneficial in such scenarios.

The capability of generating novel and similar molecules is particularly valuable in drug discovery, where exploring chemical space efficiently is crucial. As a case study, a series of 4000 derivatives of Gefitinib were generated from vicinity sampling of the latent space with a scaling factor of 0.2, resulting in 3650 chemically valid SMILES. Gefitinib is a medication used in the treatment of cancer, particularly non-small cell lung cancer (NSCLC) (*35*). It belongs to a class of drugs known as tyrosine kinase inhibitors (TKIs). Its mechanism of action involves inhibiting the activity of epidermal growth factor receptor (EGFR) tyrosine kinase (*36*). To identify potential molecular-targeted drugs with enhanced efficacy, molecular docking analysis was performed on the generated compound structures with the EGFR tyrosine kinase receptor using Schrödinger Glide (Figure 3) (*37, 38*).

**Fig. 3.**
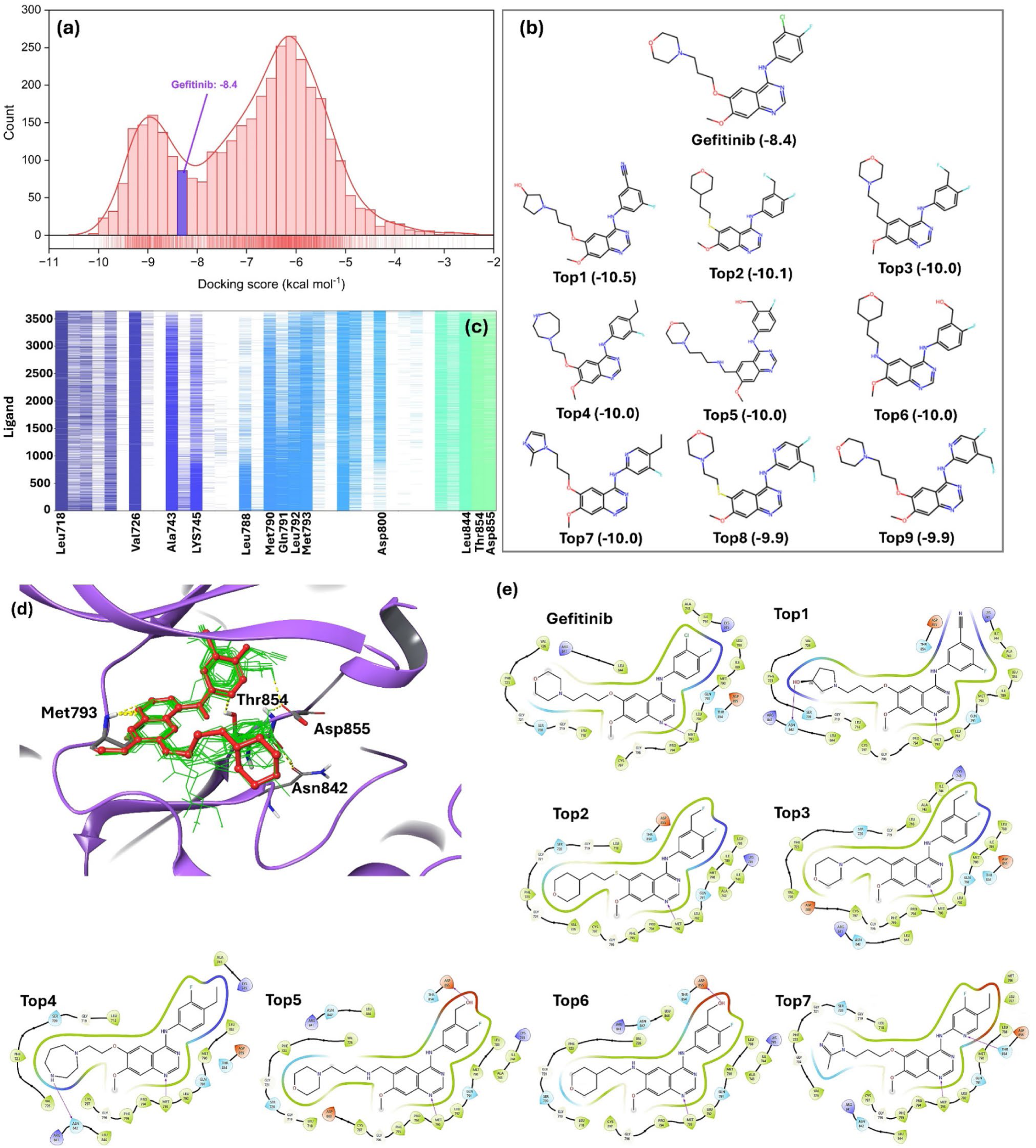
**(a)** Histogram distribution plot of docking scores for the 3650 derivatives of Gefitinib against the EGFR tyrosine kinase receptor (PDB code: 4I22) showed a bimodal distribution profile, with two peaks around −6 and −9 kcal mol^-1^. **(b)** 2D representation of Gefitinib and 9 best derivatives according to their Glide docking scores (in parenthesis; kcal mol^-1^) indicating that all the top-ranked derivatives shared the quinazoline moiety. **(c)** Interaction fingerprints for all 3650 newly generated derivatives with the target EGFR. **(d)** Binding mode of the 20 most potent derivatives (represented by green wires) and Gefitinib (illustrated in red ball and stick) within the binding site of EGFR, as predicted by the docking calculations. **(e)** 2D representation of interacting residues of the EGFR receptor within 5.0 Å radius from the ligand molecules, for Getinib and the 7 best ranked derivatives.

The exploration of novel derivatives of Gefitinib yielded multiple compounds exhibiting notably superior docking scores compared to the original compound. Intriguingly, the histogram plot of docking scores for the 3650 derivatives revealed a bimodal distribution profile, with two peaks around −6 and −9 kcal mol^-1^ (Figure 3a). Gefitinib positioned nearly midway between these peaks with a docking score of −8.4 kcal mol^-1^. Out of the newly generated derivatives, 817 exhibited docking scores surpassing that of Gefitinib, suggesting that approximately 22.4% of the derivatives generated by MolAI could potentially possess greater potency compared to Gefitinib. In a comparative study, Ochiai *et al.* (*39*) found that a natural product-oriented variational autoencoder (NP-VAE) and REINVENT (version 3.0) developed by Blaschke *et al.* (*40*) achieved similar objectives of generating novel derivatives of Gefitinib with enhanced potency. However, using the same docking protocol, their results indicated that only 15.2% and 6.8% of the new Gefitinib derivatives generated by NP-VAE and REINVENT, respectively, were predicted to be more potent than Gefitinib.

Notably, the majority of top-ranked derivatives shared the quinazoline moiety (pyrimidine-benzene fused rings) known for its significant role in EGFR interaction (Figure 3b) (*41*). Additionally, Figure 3c illustrates the confirmation of interaction fingerprints for all 3650 newly generated derivatives and the target EGFR (PDB code: 4I22) (*36*). Gefitinib and all derivatives with superior docking scores (ligands 1-1000) interacted with the backbone of MET793, crucial for hinge hydrogen bond interaction. The occurrence of this interaction diminishes for derivatives with smaller absolute docking scores (starting from ligand 1000 onwards). Similar trends were observed for multiple interactions, including Ala743, Lys745, Leu788, Met790, Gln791, Leu792, Thr854, and Aap855. Figure 3d depicts the binding mode of the Top20 best derivatives (represented by green wires) and Gefitinib (illustrated in red using ball and stick representation) within the binding site of EGFR as predicted by docking calculations. Notably, it visually demonstrates that the less solvent-exposed part of the derivatives (extending from the quinazoline moiety inward) aligns closely with Gefitinib, ensuring the formation of the critical hinge hydrogen bond with Met793. However, depending on the presence of various functional groups, derivatives could potentially establish additional interactions with other residues, such as Asn842, Thr854, and Asp855, as illustrated in Figure 3e.

The continuous latent space furthermore offers a distinctive opportunity for molecular interpolation. This allows the generation of a series of novel compounds that lie between two existing molecules by traversing the latent space from the first query molecule to the second one. If the latent vectors of the two molecules are represented as ξ_1_ and ξ_2_, respectively, the latent vector of the i^th^ generated molecule (ξ_i_) can be obtained by scanning *n* equidistant points along the path connecting these in the latent space:

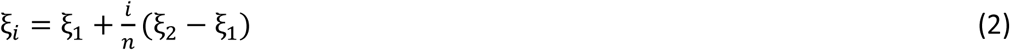

The decoder part processes these novel latent vectors and translates them into their corresponding molecules. Figure S2a showcases new compounds generated by scanning through the latent space between two query molecules. As the figure indicates, during the transition from the first query molecule towards the second one, the similarity to the first query molecule decreases, while the similarity to the second query molecule increases. The compounds situated around the halfway point in the latent space are less similar to either of the query molecules.

The interpolation can be extended beyond just two molecules and adapts any arbitrary number of molecules, *N*, using the following equation:

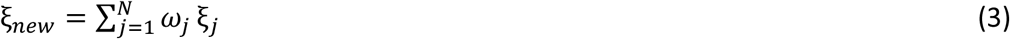

where ω_*j*_ is the weight of the latent vector of the j^th^ molecule. Any unique combination of weights that satisfies ∑_*j*=1_^*N*^ ω_*jj*_ = 1 yields a new latent vector.

As an example, we conducted a tri-molecular interpolation for the Top1, Top5, and Top7 Gefitinib derivatives generated using MolAI (Figure 3b) since besides the essential hinge hydrogen bond interaction, each of these molecules forms a new hydrogen bond interaction with the EGFR receptor through residues Asn842, Asp855, and Thr854, respectively (Figure 3e). The interpolation resulted in 54 novel molecules where 6 of them demonstrated a docking score better than −10 kcal mol^-1^ as shown in Figure S2b. The three values below each compound represents, from left to right, the weights of the latent vectors for the molecules Top1 (ω_1_), Top5 (ω_2_), and Top7 (ω_3_), respectively. The number in the parentheses indicates the corresponding docking scores (kcal mol^-1^). As Figure S2b shows, these molecules exhibit characteristics and functional groups that appear to be a blend of the attributes found in the reference molecules Top1, Top5, and Top7. However, this combination is not strictly linear. For instance, the molecule at the coordinates (0.80, 0.0, 0.20) possesses a pyrimidine-pyrrole fused ring moiety (similar to purine) rather than the quinazoline moiety present in all three reference molecules. In addition, molecules with a larger weight for a specific reference molecule (*i.e.*, points closer to one of the corners) tend to resemble that molecule more closely. For example, the molecule at the coordinate (0.20, 0.14, 0.66) bears a greater similarity to the molecule Top7 which is located at coordinate (0.00, 0.00, 1.00).

While molecular interpolation presents a promising approach for *de novo* molecular generation, it inherently restricts molecular exploration to occur within a confined elemental volume of latent space defined by query molecules. A more refined technique to thoroughly explore the entirety of the latent space involves generating semi-random latent vectors by sampling from a distribution that accurately represents the distribution of the latent space corresponding to valid molecular structures. To confirm this hypothesis, the latent vectors of 1 million unique and valid molecules from the ZINC22 database were generated and the distribution of these latent vectors were investigated. Figure S3(a) depicts the distribution of the first 16 elements of these latent vectors. Remarkably, Figure S3(a) illustrates a close adherence of the latent vector distribution to a Gaussian distribution. Hence, by deriving the parameters of each Gaussian distribution (mean and standard deviation), it becomes feasible to generate new semi-random latent vectors utilizing these distribution parameters. Following the outlined protocol, 10000 semi-random latent vectors were generated and fed into the decoder to translate them into their corresponding molecular structures. Figure S3(b) presents examples of the *de novo* generated compounds resulting from this approach. This strategy yielded a moderate proportion of valid SMILES (74%), while ensuring that the generated compounds were highly unique (98.9%). As illustrated in Figure S3(b), it is apparent that molecules featuring an amide moiety were predominantly generated, reflecting the initial distribution of the latent vectors. Indeed, from a *de novo* molecular generation perspective, this approach may not be as directly applicable as techniques like VAE, where the model learns the distribution of the latent space during training. However, it does offer a unique opportunity for sampling and generating novel compounds based on a specified distribution.

### 2.4. iLP pretraining

As described in the Methods section, the training set of the iLP model consists of approximately 10 million entries (sum over population for each output protonation state), each associated with a “state penalty” parameter. This parameter indicates the probability of a compound being in a specific protonation state, calculated using the LigPrep module in the Schrödinger software at pH = 7.0 ± 2. Figure 4a depicts the distribution of the state penalty values of the training set. In the binary classifications, the lowest protonation state of each compound was assigned to class 1, while all other protonation states were assigned to class 0. As figure 4b shows, there were approximately 7 (70%) and 3 (30%) million compounds with distinct protonation states in class 1 and class 0, respectively. Figure 4c presents the confusion matrices of the iLP binary classifier associated with 5-fold cross-validation (XV1–XV5), along with the overall confusion matrix. Figure 4d summarizes the overall binary classification metrics for the iLP model, indicating its robust performance on the validation sets. The model achieved an accuracy of 0.945 ± 0.002, a precision of 0.954 ± 0.001, a recall of 0.969 ± 0.002, and an F1-score of 0.961 ± 0.001. Additionally, the Kappa statistic and Matthew’s correlation coefficient (MCC) both stand at 0.863 ± 0.01, further indicating the model’s high reliability and predictive capability (*42*) Figures 4e and 4f provide further validation of the iLP model performance. Figure 4e displays the receiver operating characteristic (ROC) curve, with an area under the curve (AUC) of 0.99, indicating excellent model discrimination between classes. Figure 4f shows the precision-recall (PR) curve, also with an AUC of 0.99, highlighting the model’s high precision and recall in identifying true positive instances. These metrics affirm the model’s robustness and accuracy in binary classification tasks, despite the imbalanced characteristic of the training dataset with 70% class 1 and 30% class 0.

**Fig. 4.**
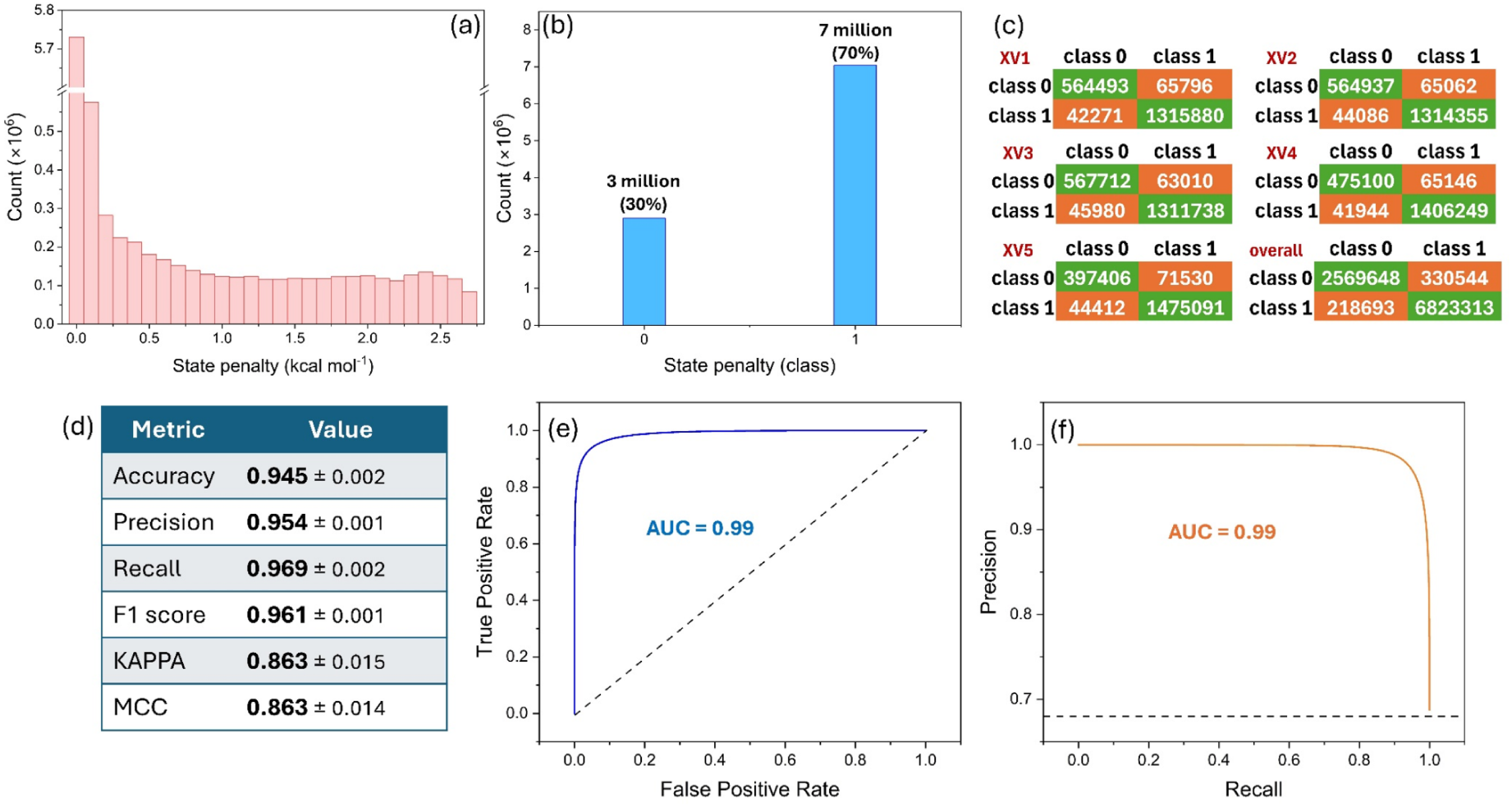
Distribution of state penalty values of the iLP’s training set **(a)** before and **(b)** after binarization. **(c)** Confusion matrices of the iLP binary classifier associated with 5-fold cross-validation (XV1–XV5), along with the overall confusion matrix. **(d)** Overall binary classification metrics for the iLP model on 5-fold cross-validation. **(e)** Receiver operating characteristic and **(f)** precision-recall curves along with the associated area under the curve (AUC) values.

### 2.5. iLP validation

As detailed in the Methods section, the performance of the pretrained iLP model was extensively investigated on various test sets. Table 1 and Figure 5a show the numbers of compounds *(#comp.*) and total number of protonation states (*#states*) along with the distribution of the classes in each test set. Figure 5a reveals that, in contrast to the training dataset (Figure 4b) that consists of 70% class 1 (correct protonation state), the test sets demonstrated totally opposite imbalanced profiles in the classes, with of only ∼10% class 1 and ∼90% class 0 (incorrect protonation states).

**Fig. 5.**
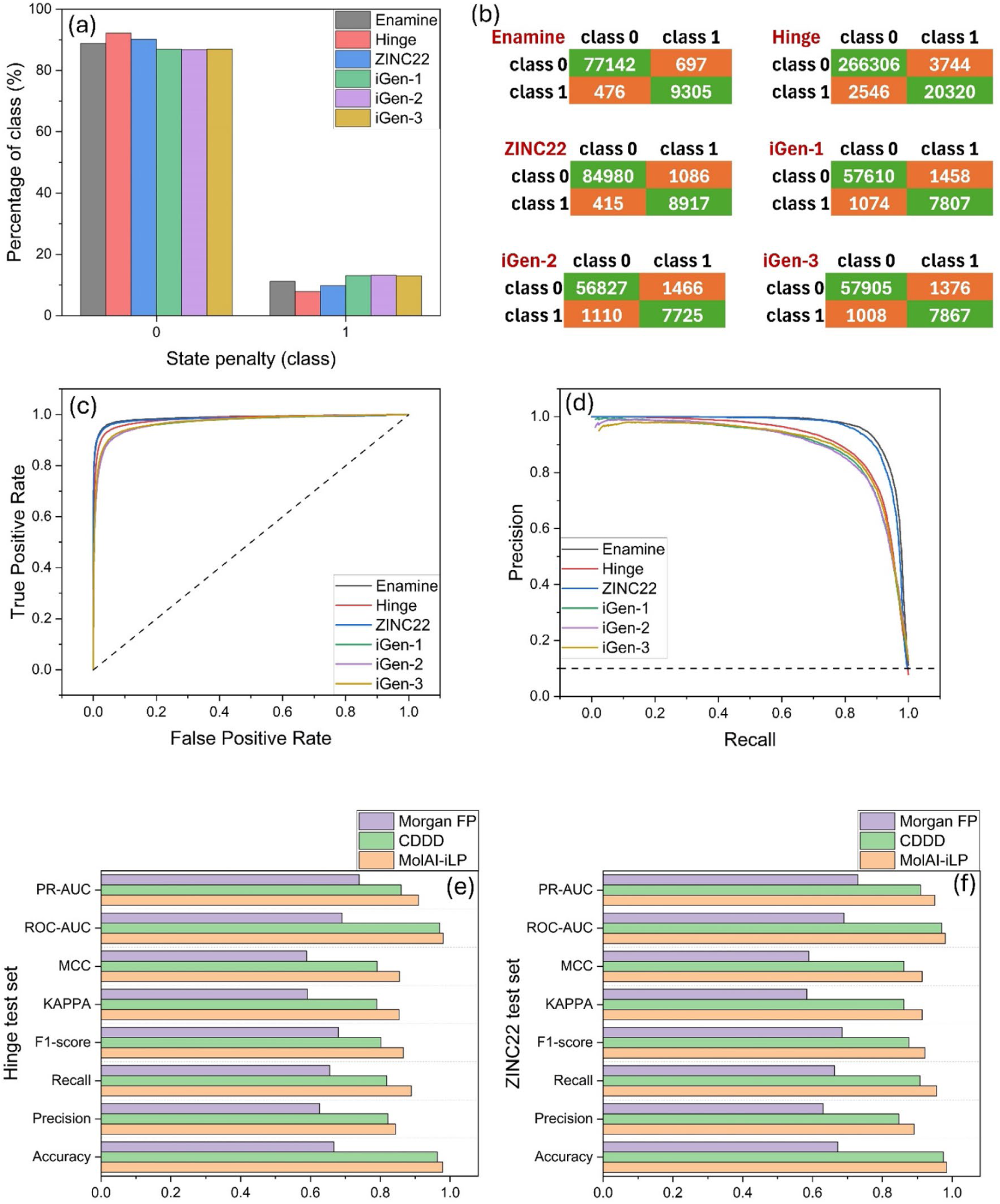
**(a)** Distribution of state penalty classes in each test set. **(b)** Confusion matrices, **(c)** ROC, and **(d)** PR curves obtained for each test set. Comparison of the performance metrics of the binary classifier obtained from Morgan FPs, original NMT autoencoder model for continuous and data-driven molecular descriptors (CDDD) by Winter *et al*. (*20*), and MolAI-driven descriptors, for **(e)** Hinge and **(f)** ZINC22 test sets.

**Table 1.**
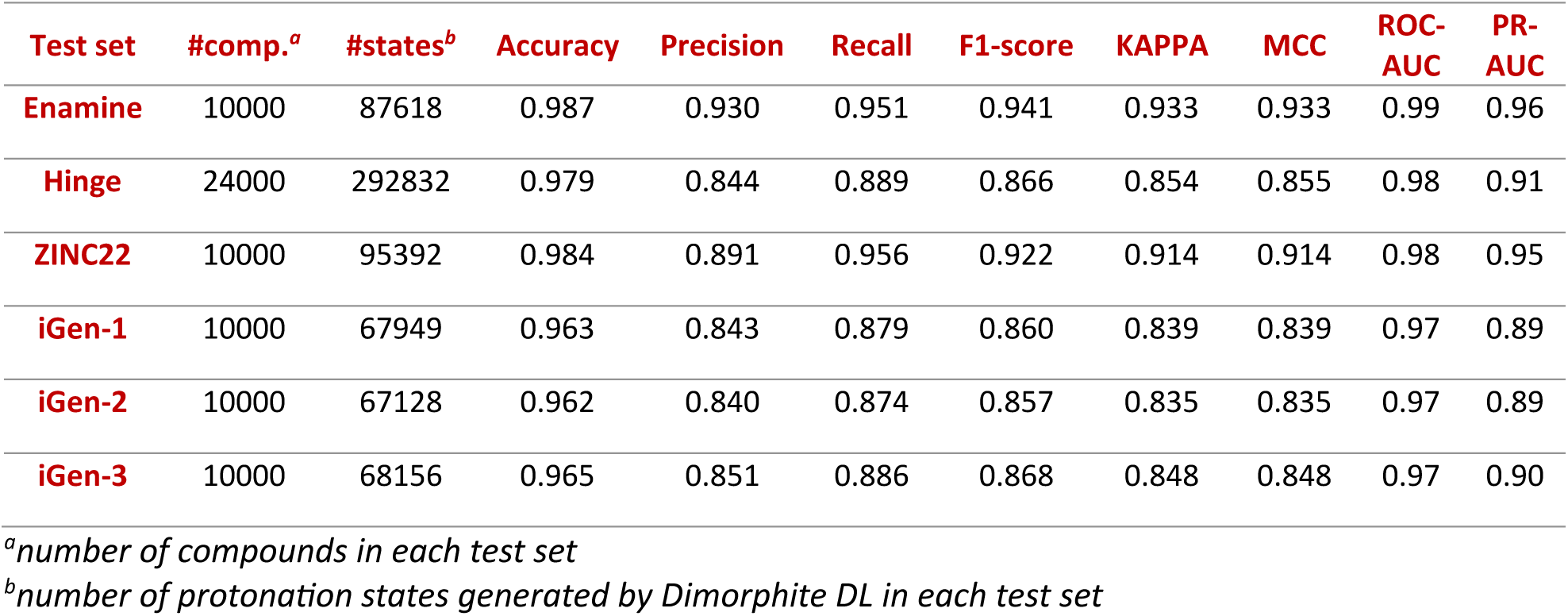
Overview of the performance metrics for the iLP model across various test sets.

| Test set | #comp.<sup>a</sup> | #states<sup>b</sup> | Accuracy | Precision | Recall | F1-score | KAPPA | MCC | ROC-AUC | PR-AUC |
| --- | --- | --- | --- | --- | --- | --- | --- | --- | --- | --- |
| Enamine | 10000 | 87618 | 0.987 | 0.930 | 0.951 | 0.941 | 0.933 | 0.933 | 0.99 | 0.96 |
| Hinge | 24000 | 292832 | 0.979 | 0.844 | 0.889 | 0.866 | 0.854 | 0.855 | 0.98 | 0.91 |
| ZINC22 | 10000 | 95392 | 0.984 | 0.891 | 0.956 | 0.922 | 0.914 | 0.914 | 0.98 | 0.95 |
| iGen-1 | 10000 | 67949 | 0.963 | 0.843 | 0.879 | 0.860 | 0.839 | 0.839 | 0.97 | 0.89 |
| iGen-2 | 10000 | 67128 | 0.962 | 0.840 | 0.874 | 0.857 | 0.835 | 0.835 | 0.97 | 0.89 |
| iGen-3 | 10000 | 68156 | 0.965 | 0.851 | 0.886 | 0.868 | 0.848 | 0.848 | 0.97 | 0.90 |

<sup>a</sup>number of compounds in each test set
<sup>b</sup>number of protonation states generated by Dimorphite DL in each test set

Table 1 provides a detailed overview of the performance metrics for the iLP model across various test sets. The binary classification metrics for the iLP model on these test sets include accuracy, precision, recall, F1-score, Kappa statistic, MCC, ROC-AUC, and PR-AUC. The iLP model shows high accuracy across all test sets, with values ranging from 0.962 to 0.987. Precision values range from 0.840 to 0.930, reflecting the proportion of true positives among the total predicted positives. Recall values and F1-score range from 0.874 to 0.956 and 0.857 to 0.941, respectively, indicating a balance between precision and recall. The Kappa statistic and MCC metric vary from 0.835 to 0.933. The ROC-AUC values, which show the model’s ability to distinguish between classes, are high, ranging from 0.97 to 0.99. Similarly, the PR-AUC values, which highlight the trade-off between precision and recall, range from 0.89 to 0.96. Figures 5b, 5c, and 5d show the confusion matrices, ROC, and PR curves obtained for each test set, respectively. Despite the inverted class distribution in the test sets compared to the training set, these metrics confirm the excellent performance of the iLP model in handling class imbalances and accurately predicting the correct protonation state of each compound.

In a large majority of successful cases, the iLP model predicts the correct protonation state of molecules with a significantly higher probability compared to other incorrect protonation states. For instance, Figure S4(a) illustrates 16 different protonation states of a molecule from the Enamine test set generated by the Dimorphite DL tool (*43*), along with the iLP-calculated probability for each protonation state being in class 1. Notably, the probability for the 8^th^ protonation state (0.9821) of this molecule is much higher than the others by a substantial margin. The next highest probability in this series is 0.2875 (11^th^ protonation state), which is considered class 0 based on a cutoff threshold of 0.5. In a few of successful cases, as shown in Figure S4(b), there are multiple protonation states classified as class 1 (probability > 0.5). However, the correct protonation state can still be identified as the one with the highest probability. This demonstrates the model’s ability to distinguish the most likely protonation state even when several states are predicted to be probable. In most unsuccessful cases, the correct protonation state is often the one with the second highest probability. Figure S4(c) illustrates such a scenario, where the highest probability (0.9967) does not correspond to the correct protonation state, but the second highest probability (0.9723) does. This highlights the iLP model’s overall effectiveness, as it often places the correct state among the top predictions even when it is not the absolute highest.

Using the same model architecture as iLP, a baseline model was trained using Morgan FPs and the prediction outcome on two test sets, Hinge and ZINC22, were compared to the original NMT autoencoder model for continuous and data-driven molecular descriptors (CDDD) by Winter *et al*. (*20*) and the MolAI-based iLP. The results are shown in Figures 5e and 5f. As these figures indicate, the CDDD model consistently followed MolAI-iLP in performance, showcasing respectable but slightly lower scores across the same metrics. In contrast, the Morgan FP model consistently showed the lowest performance among the three models, highlighting its inefficiency in this classification context. This robustness highlights MolAI’s potential utility in diverse chemical and pharmaceutical applications such as drug discovery using virtual screening and molecular docking where accurate protonation state prediction is crucial.

### 2.6. Ligand-based Virtual Screening (VS)

As detailed in the Methods section, the DUD-E database and an iterative screening protocol were used to assess the applicability of MolAI and iLP in ligand-based VS tasks. Initially, the predominant protonation state of the compounds was determined using iLP followed by the calculation of their corresponding latent vectors using MolAI. Subsequently, the maximum cosine similarity of the corresponding latent vectors to the randomly selected active set was used as the ranking criterion. The results were compared to the Tanimoto similarity index based on the Morgan fingerprints.

Figure 6a displays box plots of the average ROC-AUC values obtained by randomly selecting one, five, or ten active compounds in each iteration. The mean ROC-AUC value increases from 0.66 with one active compound to 0.80 with five, and to 0.86 with ten active compounds. The corresponding values obtained from Morgan fingerprints are 0.59, 0.70, and 0.77, respectively, as shown in Figure 6b. Figure 6c presents a box plot of the average enrichment factor at the top 1% (EF_1%_) of the ranked compounds, obtained by randomly selecting one, five, or ten active compounds. The average EF_1%_ increases from 12.1 with one active compound to 18.2 with five, and to 39.4 with ten active compounds in each iteration. Figure 6d shows the corresponding values achieved from Morgan fingerprints, which are 8.5, 14.1, and 26.2, respectively.

**Fig. 6.**
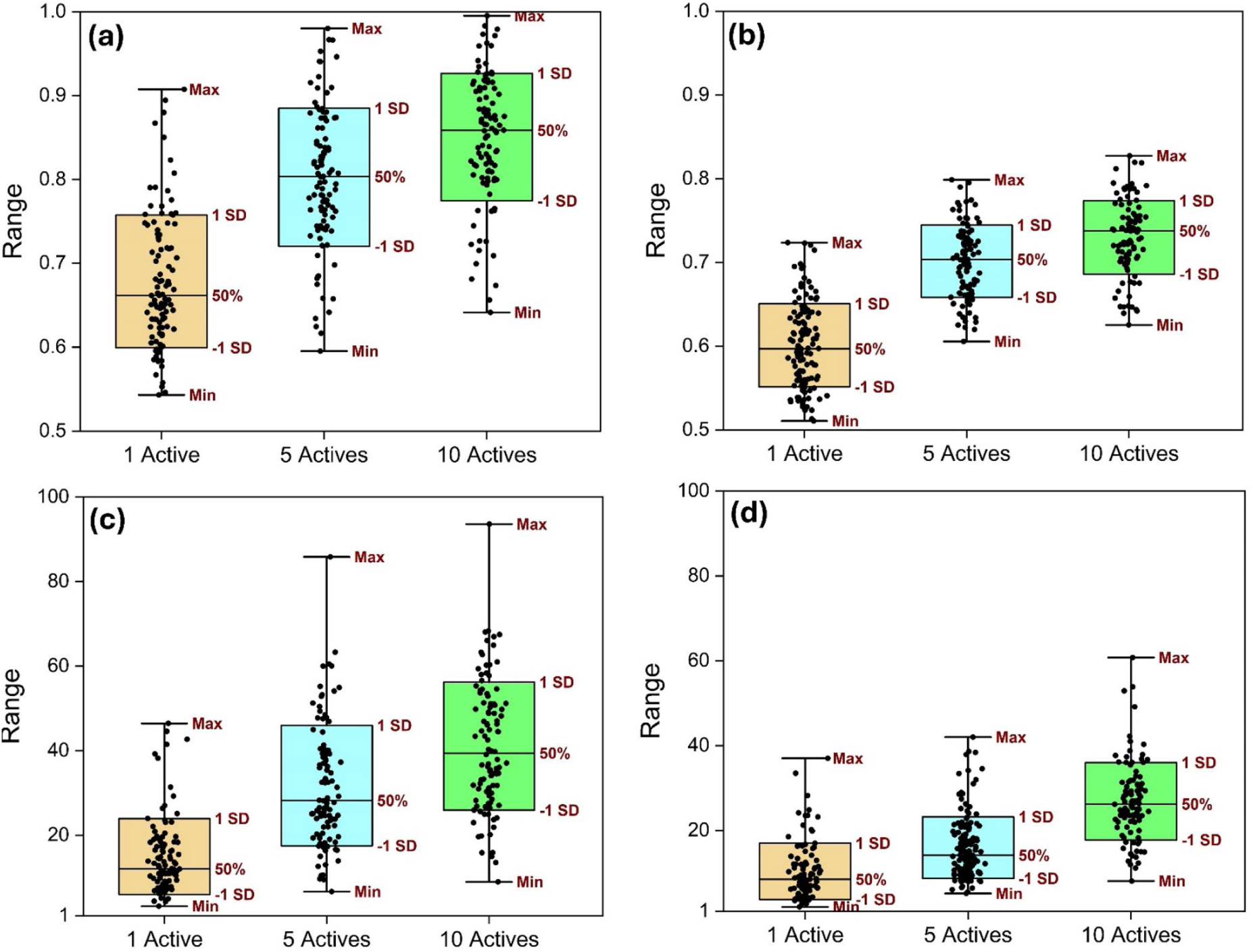
Box plots of the average ROC-AUC values from randomly selecting one, five, or ten active compounds in each iteration obtained using **(a)** MolAI descriptors and **(b)** Morgan fingerprints. Box plots of the average enrichment factor at the top 1% (EF1%) obtained using **(c)** MolAI descriptors and **(d)** Morgan fingerprints.

These results highlight the superior performance of MolAI in comparison to Morgan fingerprints, particularly in scenarios with a higher number of active compounds. The enhanced ROC-AUC values and EF_1%_ indicate that MolAI, combined with iLP, provides a more effective approach for ligand-based VS tasks.

### 2.7. iADMET pretraining

The applicability of the MolAI and iLP models was further validated by predicting 14 ADMET properties for drug-like compounds, as detailed in the Methods section under iADMET. The iADMET framework includes three regression and eleven binary classification models, as outlined in Table 2. Initially, the iLP pretrained model predicted the most predominant protonation state of the compounds in each dataset. Following this, MolAI was used to calculate the corresponding latent vectors, which were then inputted into the iADMET deep neural network for training. The performance metrics for the regression tasks included the Pearson correlation coefficient (R) and mean absolute error (MAE), while for the classification tasks, metrics such as accuracy, precision, recall, F-score, Kappa statistics, MCC, ROC-AUC, and PR-AUC were used.

Table S2 provides a summary of the performance metrics for both regression and classification datasets on the validation sets during 5-fold cross validation. For the regression tasks (caco2, lipo, sol), the models demonstrated strong performance with high Pearson correlation coefficients (R) ranging from 0.81 to 0.90 and low MAE values. This indicates that the MolAI, iLP, and iADMET models are highly effective at predicting continuous ADMET features, with the sol dataset showing particularly high predictive accuracy. Figures 7a to 7c show scatter plots of the actual and predicted values for the caco2, lipo, and sol datasets, respectively. For the classification tasks, iADMET showed robust performance across most datasets. Datasets such as bioav, bbb, hl, and herg exhibited particularly high metrics, suggesting that the models are highly reliable in correctly classifying instances in these datasets. Conversely, some datasets like cyp2c19, cyp3a4, and ames displayed relatively lower performance metrics. For instance, the cyp3a4 dataset had lower accuracy and precision compared to other datasets, indicating that the model may struggle more with this classification task. Figures 7d and 7e depict the ROC and PR plots for the classification tasks, respectively, highlighting the model’s performance in distinguishing between classes and balancing precision and recall. Figure 7f shows the corresponding confusion matrices, providing a detailed view of the model’s classification accuracy by illustrating the true positives, true negatives, false positives, and false negatives for each classification task.

**Fig. 7.**
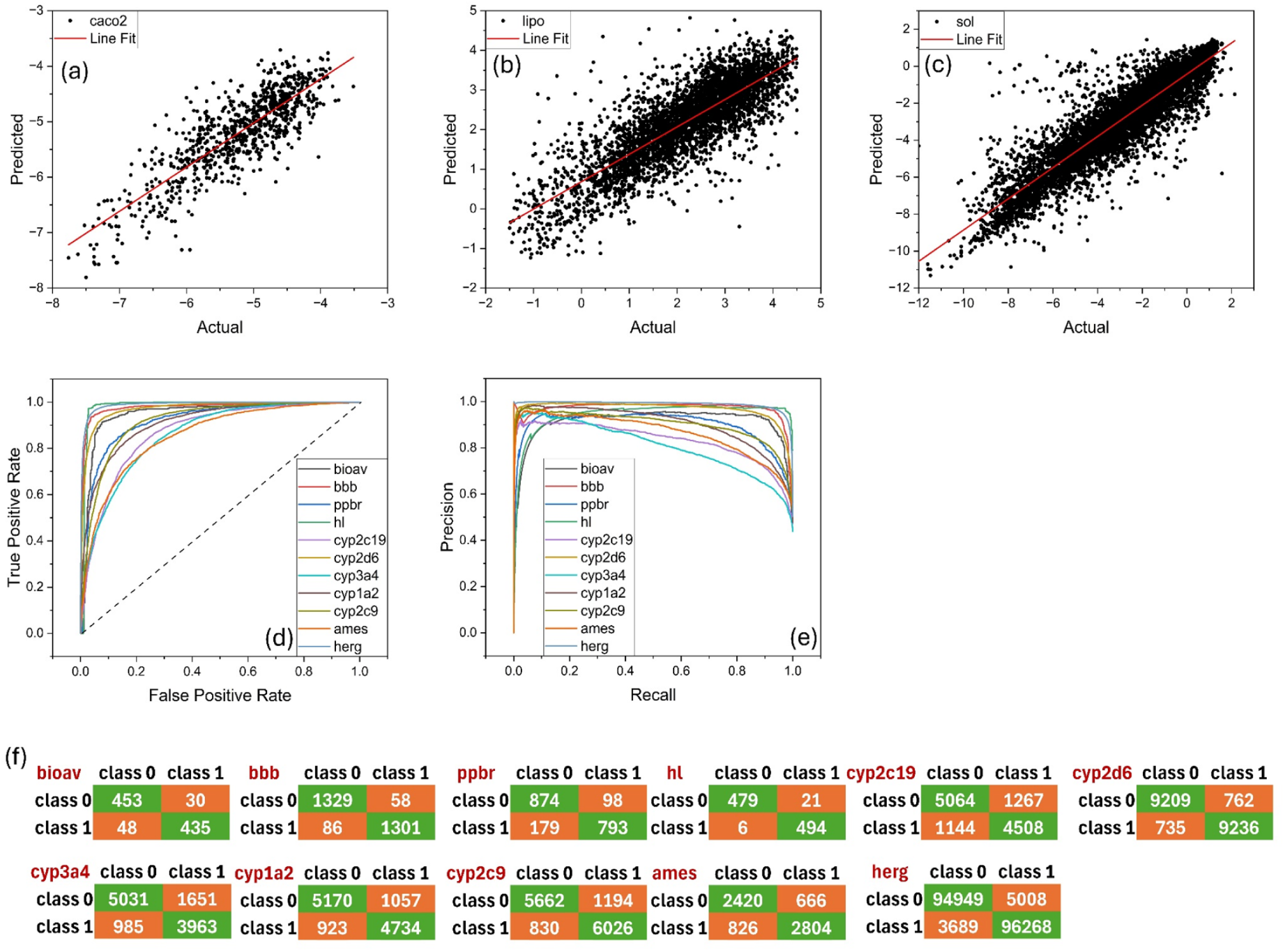
Scatter plots and best linear fit of the actual and predicted values for **(a)** caco2, **(b)** lipo, and **(c)** sol regression datasets. **(d)** ROC plots, **(e)** PR plots, and **(f)** confusion matrices for the classification datasets.

In summary, this study presents MolAI, a neural machine translation-based model designed for generating data-driven molecular descriptors. MolAI achieved 99.99% accuracy in regenerating molecules from latent vectors, proving its effectiveness in drug discovery, structure-activity relationship analysis, and *de novo* molecular generation and many more. The model outperformed traditional methods in predicting predominant protonation state of compounds at neutral pH and in ligand-based virtual screening, demonstrating superior accuracy and reliability. MolAI’s versatile descriptors also enabled the development of a robust framework) for predicting multiple ADMET features.

## 3. MATERIALS AND METHODS

### 3.1. Training databases

MolAI was trained on a huge dataset of molecular structures from the ZINC15 (*44*) and PubChem (*45*) databases. For this purpose, 200 million molecules were randomly selected from the ZINC15 and combined with the entire PubChem library (∼118 million molecules).

Subsequently, the duplicate molecules were removed and filtered using the RDKit python package (https://www.rdkit.org) with the following criteria: only organic molecules have been retained, 12 < molecular weight < 600, heavy atoms > 3, −7 < logP < 5. In addition, all counterions, solvents, and stereochemistry were removed, and the canonical SMILES representations were generated. Compounds which could not be handled by RDKit were discarded. Applying the pre-processing resulted in 221 million unique compounds. To utilize the SMILES molecular representations as input and output sequences for the translation model, start- and end-tokens were first added to the SMILES sequences. The SMILES sequences were then padded up to the size of the largest sequence in the training set (111). The padded SMILES were subsequently tokenized at the character level, except for “Br” and “Cl”, resulting in 31 unique tokens and 3 extra tokens for start, end, and pad characters. Finally, the tokenized SMILES were encoded into a one-hot vector format. Moreover, RDKit was employed to calculate 13 molecular properties for each compound: molecular weight (‘MolWt’), logP (‘MolLogP’), quantitative estimation of drug-likeness (‘QED’)(*46*), minimum and maximum partial charges (‘MinPartialCharge’ / ‘MaxPartialCharge’), number of valence electrons (‘NumValenceElectrons’), number of heavy atoms (‘HeavyAtomCount’), number of hydrogen bond acceptors and donors (‘NumHAcceptors’ / ‘NumHDonors’), Labute ASA (‘LabuteASA’) (*47*), Balaban’s J index (‘BalabanJ’) (*48*), molar refractivity (‘MolMR’) and topological polar surface area (‘TPSA’). These molecular properties were used to encourage the translation model to learn a meaningful molecular representation in terms of physiochemical features.

For iLP training, a set of 10 million random compounds were selected from the ZINC22 (*49*) database. Following the pre-processing steps mentioned above (excluding the calculation of molecular properties), this resulted in 7.6 million unique compounds, ensuring no overlap with MolAI training data. This database was prepared using the LigPrep module in the Schrödinger 2023-3 package (LigPrep, Schrödinger, LLC, New York, NY, 2023) and the Epik (*50*) tool at pH = 7.0 ± 2 using the OPLS4 (*51*) force field. The output from LigPrep comprised approximately 10 million entries (sum over population for each output protonation state), each associated with a “state penalty” parameter (energy unit) indicating the probability of a compound being in a specific protonation state. Consequently, the lowest protonation state of each compound was assigned to class 1, while all other protonation states were assigned to class 0, for the purpose of a binary classification task. The accuracy of the iLP model was extensively benchmarked against six different test sets: 10k random compounds from ZINC22, 10k random compounds from Enamine Real (https://enamine.net), 24k compounds from the hinge library of Enamine Real, and three sets of 10k AI-generated compounds using our in-house *de novo* molecular generator, iGen. Any duplicates between all benchmarking sets and the training set were first removed and canonical SMILES generated. Subsequently, up to 64 protonation states of each molecule were generated using Dimorphite DL (*43*), at pH = 7.0 ± 2. Dimorphite DL, an open-source Python library, enumerates the protonation states of small molecules, typically resulting in an output 8-10 times larger than the input size depending on the chemical composition of the molecules. In the next step, the latent vector of each compound was calculated using MolAI. The accuracy of the iLP model was then determined by comparing the predicted predominant protonation state of each compound to those obtained by LigPrep Epik.

The efficacy of the MolAI and iLP models was thoroughly evaluated for ligand-based virtual screening tasks using the Directory of Useful Decoys-Enhanced (DUD-E) database (*52*). DUD-E comprises 102 protein targets, with an average actives-to-decoys ratio of 1:50 and approximately 224 actives per target. Each decoy in the DUD-E database is a compound that has similar physicochemical properties to the active compounds but possesses a different structure. For ligand-based virtual screening, we adopted an extended version of the benchmarking strategy proposed by Riniker *et al*. (*8*) This method involves randomly selecting sets of one, five, and ten active compounds, and then ranking the remaining compounds based on the maximum cosine similarity of their corresponding latent vectors to the active set. This process was iterated 100 times with different random sets of actives, and the average ROC-AUC and enrichment factor (EF) at top 1% were calculated for each protein target.

The therapeutic data common (TDC) (*53*) database was used to train the iADMET models for ADMET feature prediction. These features include: cell effective permeability (caco2) (*54*), Bioavailability (bioav) (*55*), Lipophilicity (lipo) (*56*), Solubility (sol) (*57*), blood-brain barrier permeability (bbb) (*58*), plasma protein binding rate (ppbr) (*59*), Half Life (hl) (*60*), Inhibition of CYP P450 2C19 (cyp2c19), CYP P450 2D6 (cyp2d6), CYP P450 3A4 (cyp3a4), CYP P450 1A2 (cyp1a2), CYP P450 2C9 (cyp2c9) (*61*), Ames Mutagenicity (ames) (*62*), and hERG Central blockers (herg)(*63*). The ADMET training datasets were preprocessed using the iLP model in order to retain the predominant protonation state of the compounds. The random under-sampling technique with *random_state* = 42 and SMOTE (synthetic minority over-sampling technique) (*64*) over-sampling approach with *k_neighbors* = 5 were used to handle class imbalance in the dataset. Table S1 shows the types of prediction tasks and the number of compounds in each group of ADMET features before and after pre-processing.

### 3.2. Model Architectures

#### 3.2.1. MolAI

Figure S1 (a) shows a general overview of the MolAI architecture. For the encoding part, one-hot encoded SMILES as sequence at time step ***t*** was fed into a stack of three LSTM cells, each with 1024 memory units. The cell and hidden states of each LSTM cell were concatenated and introduced into a fully connected layer with 512 neurons and a hyperbolic tangent (‘tanh’) activation function. The output of this layer forms the latent space, which can be used as molecular descriptors during inference. The decoding part takes this latent space, re-distributes it into six separate fully connected layers, each with 1024 nodes, which are used to initialize a stack of three LSTM cells, each with 1024 memory units. The one-hot encoded SMILES at time step ***t-1*** serves as input to the first LSTM cell in the stack, utilizing professor forcing learning technique (*65*). The output of the last LSTM cell is mapped to predicted probabilities for the different tokens via a fully connected layer with 34 neurons (the size of the token library) and a *SoftMax* activation function. To enhance the model’s robustness to unseen data, noise sampling from a normal distribution with a mean of 0.0 and a standard deviation of 0.05 was applied to the latent space. The regression models consist of 13 stacks, each containing four fully connected layers with 512, 256, 128 and 1 neurons, respectively. A rectified linear unit (ReLU) activation function was used for all layers except the last one, where a linear activation function was utilized. These layers map the latent space to the molecular property vector. The *Adam* optimizer (*66*) with a learning rate of 10^−4^ was used. The batch size was set to 256. The loss function for the main translation model and each regression model was set to ‘*categorical cross entropy*’ and ‘*mean squared error*’, respectively. The full architecture of MolAI contains over 54 million trainable parameters while the encoder part consists of 21 million parameters. MolAI was trained for a total of 6 epochs (5.2 million training steps).

#### 3.2.2. iLP

The input for iLP is the latent vector (512 in length) generated by MolAI. iLP consists of a set of four fully connected layers with 1024, 512, 512 and 1 neurons, respectively, as illustrated in Figure S1(b). The ReLU activation function was used for all layers except the last, which uses a Sigmoid activation function. The *Adam* optimizer with a learning rate of 10^−4^ was used. The batch size and the loss function were set to 128 and ‘*binary cross-entropy*’, respectively. The training was conducted through 5-fold cross validation along with an early-stopping strategy with a patience of five.

#### 3.2.3. iADMET

Similar to iLP training, the input for iADMET is a latent vector generated by MolAI. iADMET comprises four fully connected layers with 2048, 1024, 1024 and 1 neurons, respectively, as shown in Figure S1(c). The ReLU activation function was utilized for all layers except the last one, which uses either a Linear or a Sigmoid activation function for the regression and classification tasks, respectively. The Adam optimizer was used with an initial learning rate of 10^−4^, which was reduced by 10% after each epoch. The loss function was set to ‘*mean absolute error*’ or ‘*binary cross-entropy*’ for the regression and classification tasks, respectively. The training was conducted through 5-fold cross validation along with an early-stopping strategy with a patience of ten.

#### 3.2.4. Hardware used

All models were trained and benchmarked using a supercomputing node equipped with 4×Nvidia A100 GPUs, 1 TB RAM and 2×Intel Xeon Gold 6338 CPU (Icelake) @2.0 GHz, totaling 64 cores, on the Alvis GPU cluster generously provided by National Academic Infrastructure for Supercomputing in Sweden (NAISS) and C3SE center for scientific and technical computing at Chalmers University of Technology in Gothenburg, Sweden.

#### 3.2.5. Timing

MolAI is capable of generating molecular latent vectors at a speed of over 4000 molecules per second on a modern NVIDIA A100 GPU, and approximately 3000 molecules per second on lower-performance GPUs such as the NVIDIA A40 and V100.

## SUPPLEMENTARY INFORMATION

Supplementary material (Figures S1-S4, Tables S1 and S2) is available for download online at at *nnnnnnnn*.

## DATA, SOFTWARE AND MATERIALS AVAILABILITY

All pretrained models—including MolAI, iLP, and iADMET—the training datasets of iADMET for re-training attempts, and associated scripts are available on the ANYO Labs GitHub repository at https://github.com/i-TripleD/MolAI-Publication.

## ACKNOWLEDGEMENTS

The authors thank the National Academic Infrastructure for Supercomputing in Sweden (NAISS) for generous allocations of computing time at supercomputing centers C3SE and NSC in part funded by The Swedish Science Research Council through grant no. 2022-06725.

## AUTHOR CONTRIBUTIONS

Conceptualization: SJM Methodology: SJM Investigation: SJM, LAE Visualization: SJM Supervision: SJM, LAE Funding acquisition: LAE Writing—original draft: SJM

Writing—review & editing: SJM, LAE

## REFERENCES

1. R. Rodríguez-Pérez, F. Miljković, J. Bajorath, Machine learning in chemoinformatics and medicinal chemistry. Annu. Rev. Biomed. Data Sci. 5, 43–65 (2022).

2. J. B. Mitchell, Machine learning methods in chemoinformatics. Wiley Interdiscip. Rev. Comput. Mol. Sci. 4, 468–481 (2014).

3. J. Vamathevan, D. Clark, P. Czodrowski, I. Dunham, E. Ferran, G. Lee, B. Li, A. Madabhushi, P. Shah, M. Spitzer, S. Zhao, Applications of machine learning in drug discovery and development. Nat. Rev. Drug Discov. 18, 463–477 (2019).

4. S. Dara, S. Dhamercherla, S. S. Jadav, C. M. Babu, M. J. Ahsan, Machine learning in drug discovery: a review. Artificial Intelligence Review 55, 1947–1999 (2022).

5. S. J. Mahdizadeh, L. A. Eriksson, iScore: A ML-Based Scoring Function for De Novo Drug Discovery. J. Chem. Inf. Model. Article ASAP. DOI: 10.1021/acs.jcim.4c02192.

6. R. Todeschini, V. Consonni, Handbook of molecular descriptors (John Wiley & Sons, 2008).

7. N. M. O’Boyle, R. A. Sayle, Comparing structural fingerprints using a literature-based similarity benchmark. J. Cheminform. 8, 1–14 (2016).

8. S. Riniker, G. A. Landrum, Open-source platform to benchmark fingerprints for ligand-based virtual screening. J. Cheminform. 5, 26 (2013).

9. S. Jaeger, S. Fulle, S. Turk, Mol2vec: unsupervised machine learning approach with chemical intuition. J. Chem. Inf. Model. 58, 27–35 (2018).

10. Y. LeCun, Y. Bengio, G. Hinton, Deep learning. Nature 521, 436–444 (2015).

11. I. Goodfellow, Y. Bengio, A. Courville, Deep learning (MIT press, 2016).

12. S. Kearnes, K. McCloskey, M. Berndl, V. Pande, P. Riley, Molecular graph convolutions: moving beyond fingerprints. J. Comput. Aided Mol. Des. 30, 595–608 (2016).

13. D. Weininger, SMILES, a chemical language and information system. 1. Introduction to methodology and encoding rules. J. Chem. Inf. Comput. Sci. 28, 31–36 (1988).

14. K. W. Church, Word2Vec. Nat. Lang. Eng. 23, 155–162 (2017).

15. R. Gómez-Bombarelli, J. N. Wei, D. Duvenaud, J. M. Hernández-Lobato, B. Sánchez-Lengeling, D. Sheberla, J. Aguilera-Iparraguirre, T. D. Hirzel, R. P. Adams, A. Aspuru-Guzik, Automatic chemical design using a data-driven continuous representation of molecules. ACS Cent. Sci. 4, 268–276 (2018).

16. D. P. Kingma, M. Welling, Auto-encoding variational bayes. arXiv preprint arXiv:1312.6114, (2013).

17. J. Gu, Z. Wang, J. Kuen, L. Ma, A. Shahroudy, B. Shuai, T. Liu, X. Wang, G. Wang, J. Cai, Recent advances in convolutional neural networks. Pattern recognit. 77, 354–377 (2018).

18. K. Cho, B. Van Merriënboer, C. Gulcehre, D. Bahdanau, F. Bougares, H. Schwenk, Y. Bengio, Learning phrase representations using RNN encoder-decoder for statistical machine translation. arXiv preprint arXiv:1406.1078, (2014).

19. I. Sutskever, O. Vinyals, Q. V. Le, Sequence to sequence learning with neural networks. Adv. Neural Inf. Process. Syst. 27, (2014).

20. R. Winter, F. Montanari, F. Noé, D. Clevert, Learning continuous and data-driven molecular descriptors by translating equivalent chemical representations. Chem. Sci., 10, 1692–1701 (2019).

21. D. Bahdanau, K. Cho, Y. Bengio, Neural machine translation by jointly learning to align and translate. arXiv preprint arXiv:1409.0473, (2014).

22. K. Greff, R. K. Srivastava, J. Koutník, B. R. Steunebrink, J. Schmidhuber, LSTM: A search space odyssey. IEEE Trans. Neural Netw. Learn. Syst. 28, 2222–2232 (2016).

23. R. Cahuantzi, X. Chen, S. Güttel, A Comparison of LSTM and GRU Networks for Learning Symbolic Sequences. (Springer, 2023), pp. 771–785.

24. M. S. Park, C. Gao, H. A. Stern, Estimating binding affinities by docking/scoring methods using variable protonation states. Proteins 79, 304–314 (2011).

25. T. Ten Brink, T. E. Exner, pK a based protonation states and microspecies for protein– ligand docking. J. Comput. Aided Mol. Des. 24, 935–942 (2010).

26. N. Brooijmans, C. Humblet, Chemical space sampling by different scoring functions and crystal structures. J. Comput. Aided Mol. Des. 24, 433–447 (2010).

27. F. Milletti, A. Vulpetti, Tautomer preference in PDB complexes and its impact on structure-based drug discovery. J. Chem. Inf. Model. 50, 1062–1074 (2010).

28. E. Yuriev, P. A. Ramsland, Latest developments in molecular docking: 2010–2011 in review. J. Mol. Recognit. 26, 215–239 (2013).

29. D. Chen, N. Oezguen, P. Urvil, C. Ferguson, S. M. Dann, T. C. Savidge, Regulation of protein-ligand binding affinity by hydrogen bond pairing. Sci. Adv. 2, e1501240 (2016).

30. H. Van De Waterbeemd, E. Gifford, ADMET in silico modelling: towards prediction paradise? Nat. Rev. Drug Discov. 2, 192–204 (2003).

31. P. Atallah, K. B. Wagener, M. D. Schulz, ADMET: The future revealed. Macromolecules 46, 4735–4741 (2013).

32. S. H. Bertz, The first general index of molecular complexity. J. Am. Chem. Soc. 103, 3599–3601 (1981).

33. P. Ertl, A. Schuffenhauer, Estimation of synthetic accessibility score of drug-like molecules based on molecular complexity and fragment contributions. J. Cheminform. 1, 1–11 (2009).

34. D. Bajusz, A. Rácz, K. Héberger, Why is Tanimoto index an appropriate choice for fingerprint-based similarity calculations? J. Cheminform. 7, 1–13 (2015).

35. R. S. Herbst, M. Fukuoka, J. Baselga, Gefitinib—a novel targeted approach to treating cancer. Nat. Rev. Cancer 4, 956–965 (2004).

36. K. S. Gajiwala, J. Feng, R. Ferre, K. Ryan, O. Brodsky, S. Weinrich, J. C. Kath, A. Stewart, Insights into the aberrant activity of mutant EGFR kinase domain and drug recognition. Structure 21, 209–219 (2013).

37. R. A. Friesner, J. L. Banks, R. B. Murphy, T. A. Halgren, J. J. Klicic, D. T. Mainz, M. P. Repasky, E. H. Knoll, M. Shelley, J. K. Perry, Glide: a new approach for rapid, accurate docking and scoring. 1. Method and assessment of docking accuracy. J. Med. Chem. 47, 1739–1749 (2004).

38. T. A. Halgren, R. B. Murphy, R. A. Friesner, H. S. Beard, L. L. Frye, W. T. Pollard, J. L. Banks, Glide: a new approach for rapid, accurate docking and scoring. 2. Enrichment factors in database screening. J. Med. Chem. 47, 1750–1759 (2004).

39. T. Ochiai, T. Inukai, M. Akiyama, K. Furui, M. Ohue, N. Matsumori, S. Inuki, M. Uesugi, T. Sunazuka, K. Kikuchi, Variational autoencoder-based chemical latent space for large molecular structures with 3D complexity. Commun. Chem. 6, 249 (2023).

40. T. Blaschke, J. Arús-Pous, H. Chen, C. Margreitter, C. Tyrchan, O. Engkvist, K. Papadopoulos, A. Patronov, REINVENT 2.0: an AI tool for de novo drug design. J. Chem. Inf. Model. 60, 5918–5922 (2020).

41. D. A. Cross, S. E. Ashton, S. Ghiorghiu, C. Eberlein, C. A. Nebhan, P. J. Spitzler, J. P. Orme, M. R. V. Finlay, R. A. Ward, M. J. Mellor, AZD9291, an irreversible EGFR TKI, overcomes T790M-mediated resistance to EGFR inhibitors in lung cancer. Cancer Discov. 4, 1046–1061 (2014).

42. Ž. Vujović, Classification model evaluation metrics. Int. J. Adv. Comput. Sci. Appl. 12, 599–606 (2021).

43. P. J. Ropp, J. C. Kaminsky, S. Yablonski, J. D. Durrant, Dimorphite-DL: an open-source program for enumerating the ionization states of drug-like small molecules. J. Cheminform. 11, 1–8 (2019).

44. T. Sterling, J. J. Irwin, ZINC 15–ligand discovery for everyone. J. Chem. Inf. Model. 55, 2324–2337 (2015).

45. S. Kim, J. Chen, T. Cheng, A. Gindulyte, J. He, S. He, Q. Li, B. A. Shoemaker, P. A. Thiessen, B. Yu, PubChem 2023 update. Nucleic Acids Res. 51, D1373–D1380 (2023).

46. G. R. Bickerton, G. V. Paolini, J. Besnard, S. Muresan, A. L. Hopkins, Quantifying the chemical beauty of drugs. Nat. Chem. 4, 90–98 (2012).

47. P. Labute, A widely applicable set of descriptors. J. Mol. Graph. Model. 18, 464–477 (2000).

48. A. T. Balaban, Highly discriminating distance-based topological index. Chem. Phys. Lett. 89, 399–404 (1982).

49. B. I. Tingle, K. G. Tang, M. Castanon, J. J. Gutierrez, M. Khurelbaatar, C. Dandarchuluun, Y. S. Moroz, J. J. Irwin, ZINC-22─ A free multi-billion-scale database of tangible compounds for ligand discovery. J. Chem. Inf. Model. 63, 1166–1176 (2023).

50. J. C. Shelley, A. Cholleti, L. L. Frye, J. R. Greenwood, M. R. Timlin, M. Uchimaya, Epik: a software program for pK a prediction and protonation state generation for drug-like molecules. J. Comput. Aided Mol. Des. 21, 681–691 (2007).

51. C. Lu, C. Wu, D. Ghoreishi, W. Chen, L. Wang, W. Damm, G. A. Ross, M. K. Dahlgren, E. Russell, C. D. Von Bargen, OPLS4: Improving force field accuracy on challenging regimes of chemical space. J. Chem. Theory Comput. 17, 4291–4300 (2021).

52. M. M. Mysinger, M. Carchia, J. J. Irwin, B. K. Shoichet, Directory of useful decoys, enhanced (DUD-E): better ligands and decoys for better benchmarking. J. Med. Chem. 55, 6582–6594 (2012).

53. K. Huang, T. Fu, W. Gao, Y. Zhao, Y. Roohani, J. Leskovec, C. W. Coley, C. Xiao, J. Sun, M. Zitnik, Artificial intelligence foundation for therapeutic science. Nat. Chem. Biol. 18, 1033–1036 (2022).

54. N.-N. Wang, J. Dong, Y.-H. Deng, M.-F. Zhu, M. Wen, Z.-J. Yao, A.-P. Lu, J.-B. Wang, D.-S. Cao, ADME properties evaluation in drug discovery: prediction of Caco-2 cell permeability using a combination of NSGA-II and boosting. J. Chem. Inf. Model. 56, 763–773 (2016).

55. C.-Y. Ma, S.-Y. Yang, H. Zhang, M.-L. Xiang, Q. Huang, Y.-Q. Wei, Prediction models of human plasma protein binding rate and oral bioavailability derived by using GA–CG– SVM method. J. Pharm. Biomed. Anal. 47, 677–682 (2008).

56. Z. Wu, B. Ramsundar, E. N. Feinberg, J. Gomes, C. Geniesse, A. S. Pappu, K. Leswing, V. Pande, MoleculeNet: a benchmark for molecular machine learning. Chem. Sci. 9, 513–530 (2018).

57. M. C. Sorkun, A. Khetan, S. Er, AqSolDB, a curated reference set of aqueous solubility and 2D descriptors for a diverse set of compounds. Sci. Data 6, 143 (2019).

58. I. F. Martins, A. L. Teixeira, L. Pinheiro, A. O. Falcao, A Bayesian approach to in silico blood-brain barrier penetration modeling. J. Chem. Inf. Model. 52, 1686–1697 (2012).

59. M. Wenlock, N. Tomkinson, Experimental in vitro DMPK and physicochemical data on a set of publicly disclosed compounds. (EMBL, 2015).

60. R. S. Obach, F. Lombardo, N. J. Waters, Trend analysis of a database of intravenous pharmacokinetic parameters in humans for 670 drug compounds. Drug Metab. Disposition 36, 1385–1405 (2008).

61. H. Veith, N. Southall, R. Huang, T. James, D. Fayne, N. Artemenko, M. Shen, J. Inglese, C. P. Austin, D. G. Lloyd, Comprehensive characterization of cytochrome P450 isozyme selectivity across chemical libraries. Nat. Biotechnol. 27, 1050–1055 (2009).

62. C. Xu, F. Cheng, L. Chen, Z. Du, W. Li, G. Liu, P. W. Lee, Y. Tang, In silico prediction of chemical Ames mutagenicity. J. Chem. Inf. Model. 52, 2840–2847 (2012).

63. F. Du, H. Yu, B. Zou, J. Babcock, S. Long, M. Li, hERGCentral: a large database to store, retrieve, and analyze compound-human Ether-a-go-go related gene channel interactions to facilitate cardiotoxicity assessment in drug development. Assay Drug Dev. Technol. 9, 580–588 (2011).

64. N. V. Chawla, K. W. Bowyer, L. O. Hall, W. P. Kegelmeyer, SMOTE: synthetic minority over-sampling technique. J. Artif. Intell. Res. 16, 321–357 (2002).

65. A. Goyal, A. M. Lamb, Y. Zhang, S. Zhang, A. Courville, Y. Bengio, Professor forcing: A new algorithm for training recurrent networks. Adv. Neural Inf. Process. Syst. 29, 1–9 (2016).

66. D. P. Kingma, J. Ba, Adam: A method for stochastic optimization. arXiv preprint arXiv:1412.6980, (2014).

