## Supplementary Figures S1-S4 and Tables S1 - S2. for "MolAI: A Deep Learning Framework for Data-driven Molecular Descriptor Generation and Advanced Drug Discovery Applications"

**This PDF file includes:**

Figs. S1 to S4

Tables S1 and S2

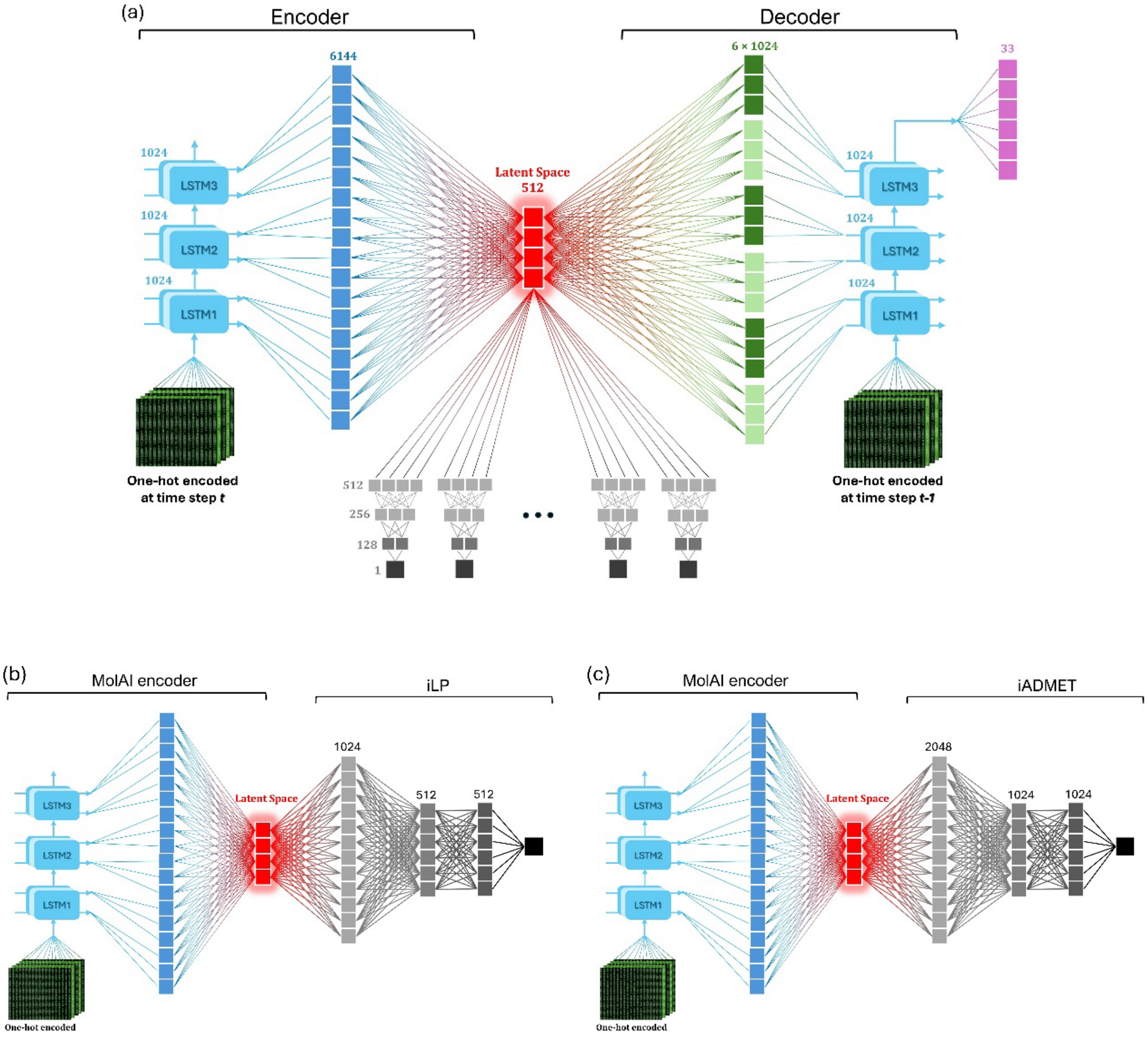


**Fig. S1.** The model architecture for **(a)** MolAI, **(b)** iLP, and **(c)** iADMET.

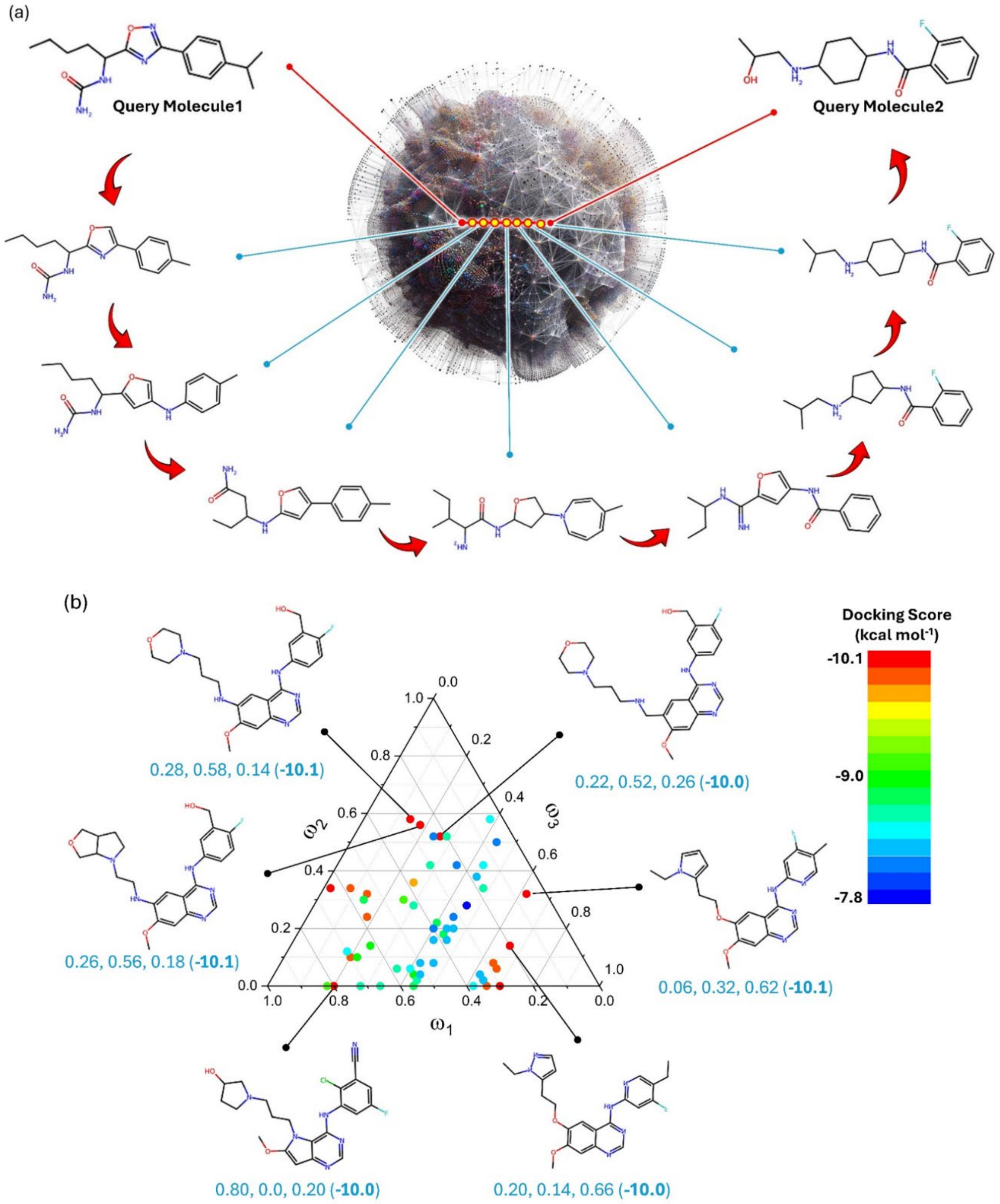


**Fig. S2. (a)** Example of new compounds generated using bi-molecular interpolation by scanning through the latent space between two query molecules. **(b)** A tri-molecular interpolation for Top1, Top5, and Top,7 Gefitinib derivatives generated using MolAI resulted in 54 novel molecules where 6 of them demonstrated a docking score better than -10 kcal mol$^{-1}$. The three values below each compound represents, from left to right, the weights of the latent vectors for the molecules Top1 ($\omega_1$), Top5 ($\omega_2$), and Top7 ($\omega_3$), respectively. The numbers in the parentheses indicate the corresponding Glide docking scores (kcal mol$^{-1}$).

(a)

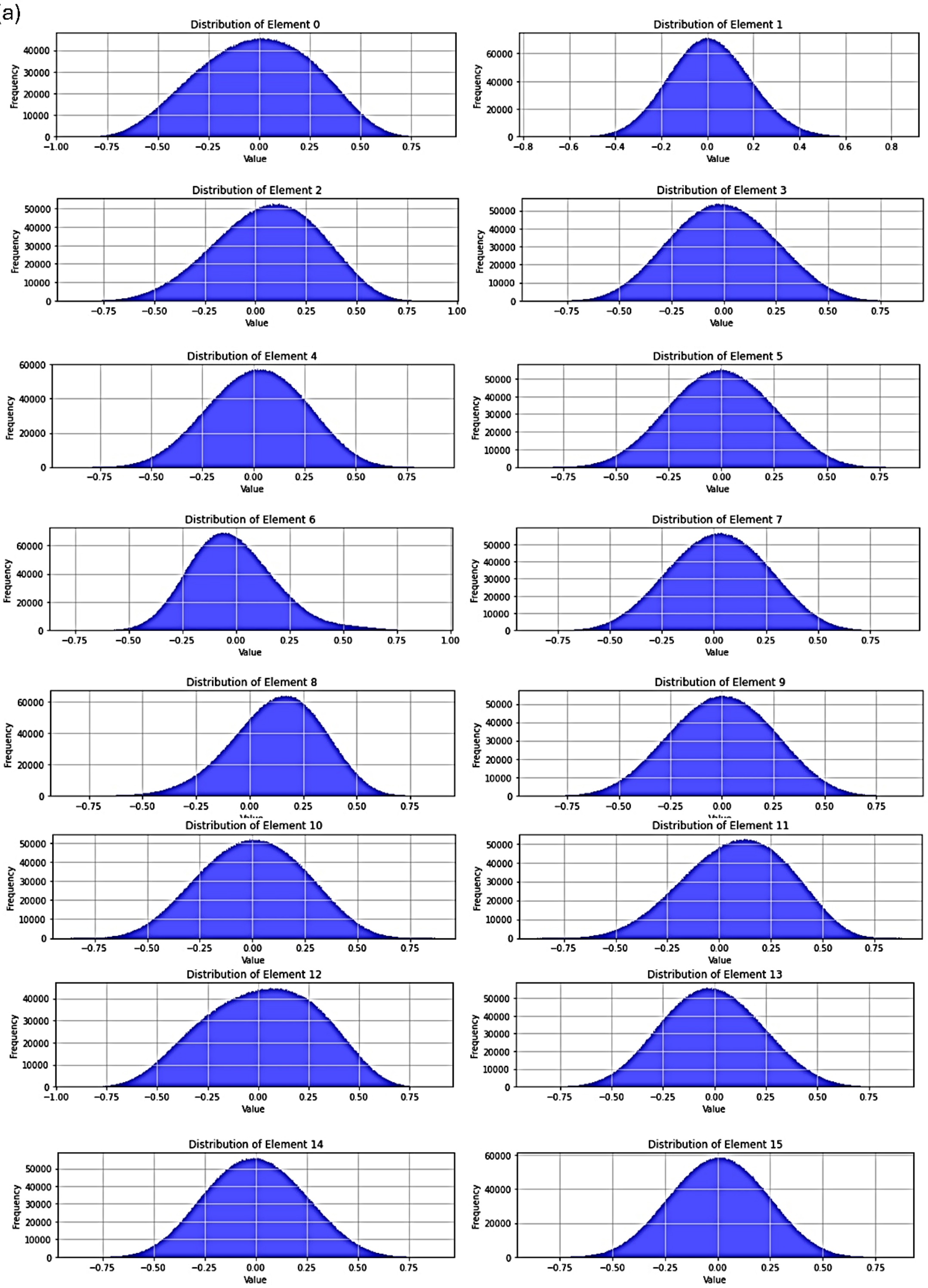
Distribution of Element 0
Distribution of Element 1
Distribution of Element 2
Distribution of Element 3
Distribution of Element 4
Distribution of Element 5
Distribution of Element 6
Distribution of Element 7
Distribution of Element 8
Distribution of Element 9
Distribution of Element 10
Distribution of Element 11
Distribution of Element 12
Distribution of Element 13
Distribution of Element 14
Distribution of Element 15
Frequency
Value

(b)

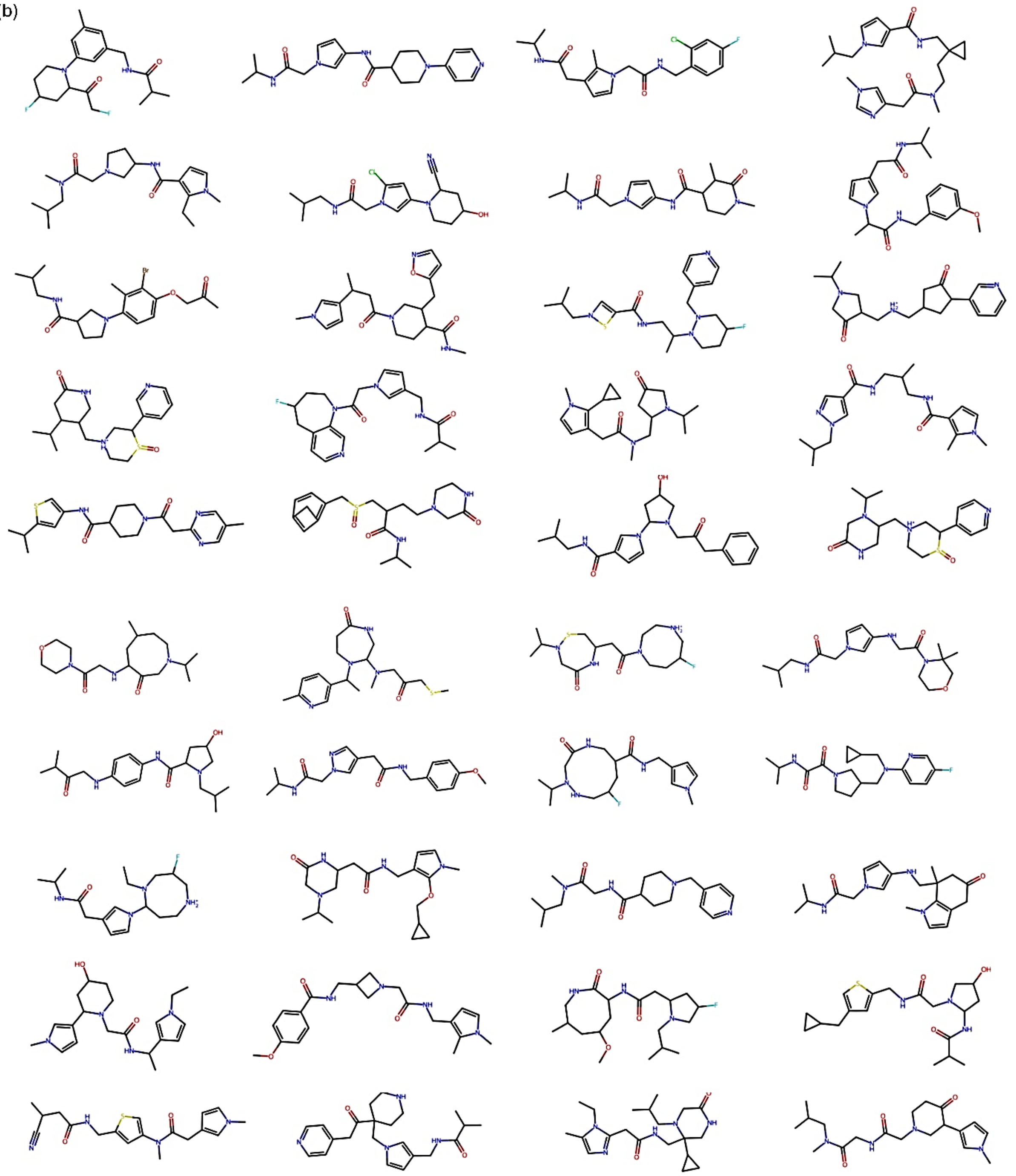

**Fig. S3. (a)** Distribution of the first 16 elements of latent vectors generated by MolAI for 1 million unique and valid molecules from the ZINC22 database. **(b)** Examples of de. novo generated compounds resulting from random sampling based on the aforementioned latent space distributions.

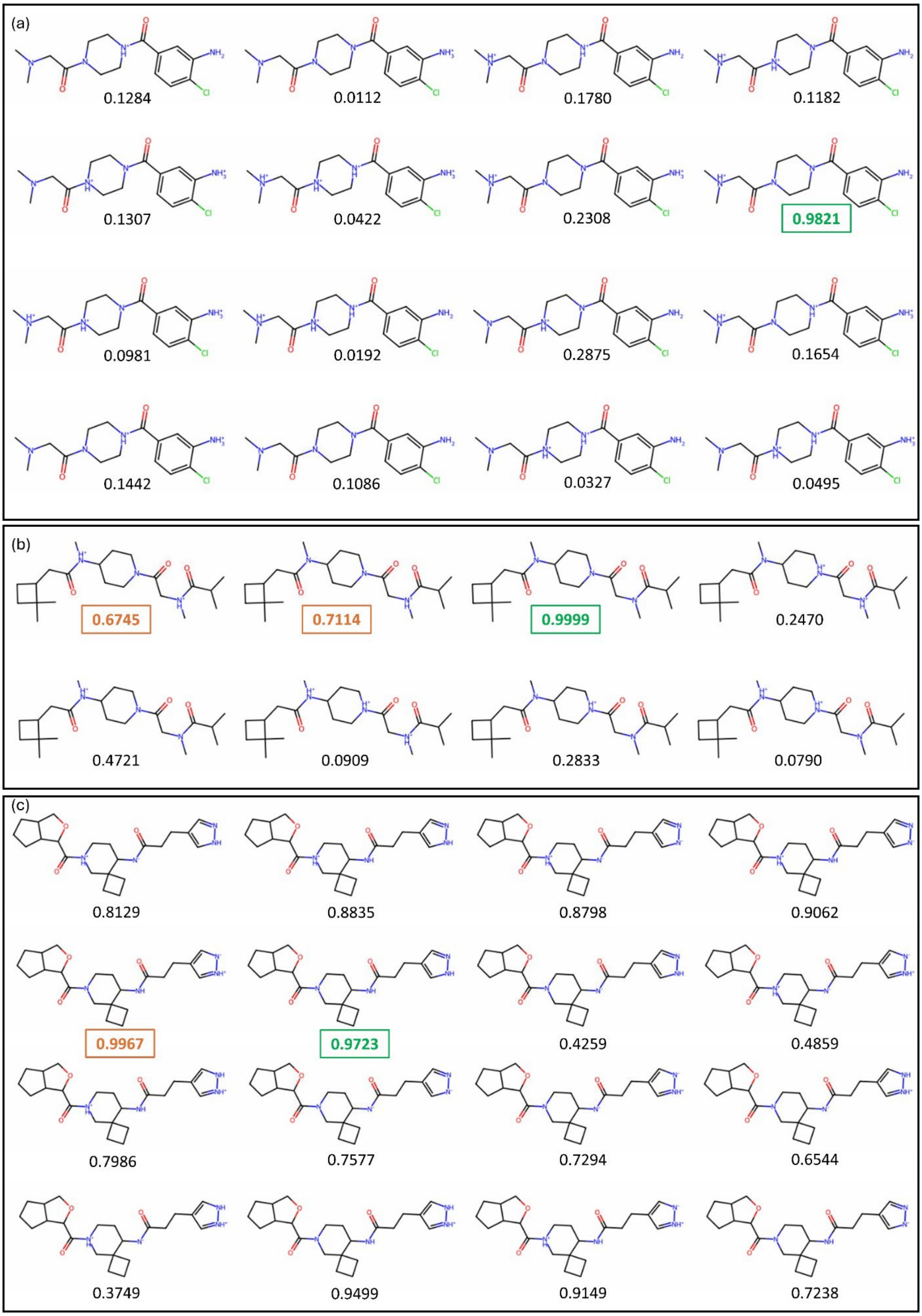


**Fig. S4. (a)** 16 different protonation states of a molecule from the Enamine test set generated by the Dimorphite DL tool (Ref.09.in.main.text), along with the iLP-calculated probability for each protonation state being in class 1. The iLP model predicts the correct protonation state (highlighted in green) with a significantly higher probability compared to other incorrect protonation states. **(b)** Multiple protonation states classified as class 1 (probability > 0.5, highlighted in orange) but the correct protonation state can still be identified as the one with the highest probability. **(c)** In most unsuccessful cases, the correct protonation state is often the one with the second highest probability (highlighted in green).

**Table S1.** Fourteen datasets used for benchmarking MolAI-driven molecular descriptors through the training of the iADMET framework for ADMET features predictions.

| Dataset | Acronym | Task | Datapoints before pre-processing | Datapoints after pre-processing | Definition of class 1 |
|---|---|---|---|---|---|
| **Cell Effective Permeability** | caco2 | Regression | 906 | 865 | - |
| **Lipophilicity** | lipo | Regression | 4200 | 4174 | - |
| **Solubility** | sol | Regression | 9982 | 8217 | - |
| **Bioavailability** | bioav* | Classification | 640 | 966 | active |
| **Blood-Brain Barrier Perm.** | bbb* | Classification | 1975 | 2774 | permeable |
| **Plasma Protein Binding Rate** | ppbr* | Classification | 1614 | 1944 | > 90% |
| **Half Life** | hl* | Classification | 667 | 1000 | >24h |
| **CYP P450 2C19** | cyp2c19 | Classification | 12665 | 11982 | inhibition |
| **CYP P450 2D6** | cyp2d6* | Classification | 13130 | 19942 | inhibition |
| **CYP P450 3A4** | cyp3a4 | Classification | 12328 | 11630 | inhibition |
| **CYP P450 1A2** | cyp1a2 | Classification | 12579 | 11884 | inhibition |
| **CYP P450 2C9** | cyp2c9* | Classification | 12092 | 13712 | inhibition |
| **Ames Mutagenicity** | ames | Classification | 7255 | 6717 | mutagenic |
| **hERG Central Blockers** | herg* | Classification | 306893 | 199914 | blocker |

**The SMOTE method was utilized to handle class imbalance*

**Table S2.** Summary of the iADMET framework performance metrics for regression and classification datasets on the validation sets using 5-fold cross validation

| Dataset | R[a] | MAE[b] | Accuracy | Precision | Recall | F1-score | KAPPA | MCC | ROC-AUC | PR-AUC |
|---|---|---|---|---|---|---|---|---|---|---|
| caco2 | **0.833** ±0.04 | **0.340** ±0.03 | - | - | - | - | - | - | - | - |
| lipo | **0.813** ±0.02 | **0.516** ±0.02 | - | - | - | - | - | - | - | - |
| sol | **0.901** ±0.01 | **0.640** ±0.01 | - | - | - | - | - | - | - | - |
| bioav | - | - | **0.919** ±0.011 | **0.935** ±0.006 | **0.900±** 0.027 | **0.917** ±0.016 | **0.837** ±0.023 | **0.838** ±0.022 | **0.95** | **0.91** |
| bbb | - | - | **0.948** ±0.012 | **0.957** ±0.002 | **0.938±** 0.023 | **0.947** ±0.012 | **0.896** ±0.024 | **0.896** ±0.024 | **0.98** | **0.97** |
| ppbr | - | - | **0.858** ±0.018 | **0.892** ±0.030 | **0.816±** 0.057 | **0.850** ±0.026 | **0.715** ±0.037 | **0.720** ±0.033 | **0.92** | **0.89** |
| hl | - | - | **0.973** ±0.012 | **0.959** ±0.024 | **0.988±** 0.004 | **0.973** ±0.013 | **0.946** ±0.025 | **0.947** ±0.024 | **0.98** | **0.94** |
| cyp2c19 | - | - | **0.799** ±0.011 | **0.781** ±0.009 | **0.798±** 0.035 | **0.789** ±0.015 | **0.597** ±0.022 | **0.597** ±0.022 | **0.87** | **0.83** |
| cyp2d6 | - | - | **0.925** ±0.001 | **0.924** ±0.009 | **0.926±** 0.012 | **0.925** ±0.002 | **0.850** ±0.002 | **0.850** ±0.002 | **0.97** | **0.97** |
| cyp3a4 | - | - | **0.733** ±0.013 | **0.708** ±0.032 | **0.801±** 0.028 | **0.751** ±0.008 | **0.545** ±0.023 | **0.550** ±0.021 | **0.86** | **0.81** |
| cyp1a2 | - | - | **0.833** ±0.005 | **0.818** ±0.014 | **0.836±** 0.030 | **0.856** ±0.008 | **0.705** ±0.018 | **0.707** ±0.017 | **0.92** | **0.90** |
| cyp2c9 | - | - | **0.852** ±0.009 | **0.836** ±0.0027 | **0.879±** 0.041 | **0.714** ±0.020 | **0.565** ±0.025 | **0.566** ±0.025 | **0.87** | **0.76** |
| ames | - | - | **0.778** ±0.020 | **0.808** ±0.022 | **0.772±** 0.021 | **0.790** ±0.019 | **0.554** ±0.040 | **0.555** ±0.040 | **0.86** | **0.86** |
| herg | - | - | **0.956** ±0.005 | **0.951** ±0.010 | **0.963±** 0.010 | **0.957** ±0.005 | **0.913** ±0.009 | **0.913** ±0.009 | **0.99** | **0.99** |

[a]Pearson.correlation.coefficient

[b]mean.absolute.error
